# Comparing synaptic proteomes across seven mouse models for autism reveals molecular subtypes and deficits in Rho GTPase signaling

**DOI:** 10.1101/2021.02.02.429412

**Authors:** Abigail U. Carbonell, Carmen Freire-Cobo, Ilana V. Deyneko, Hediye Erdjument-Bromage, Amy E. Clipperton-Allen, Randall L. Rasmusson, Damon T. Page, Thomas A. Neubert, Bryen A. Jordan

**Affiliations:** Dominick P. Purpura Department of Neuroscience, Albert Einstein College of Medicine, Bronx, NY, USA; Department of Psychiatry and Behavioral Sciences, Albert Einstein College of Medicine, Bronx, NY, USA; Department of Cell Biology and Kimmel Center for Biology and Medicine at the Skirball Institute, New York University School of Medicine, New York, NY, USA; Department of Neuroscience, The Scripps Research Institute Florida, Jupiter, FL, USA; Department of Physiology and Biophysics, Jacobs School of Medicine & Biomedical Sciences, University at Buffalo, Buffalo, NY, USA

**Author notes:** **Corresponding author and lead contact:** Bryen A. Jordan, Ph.D., Dominick P. Purpura Department of Neuroscience, Albert Einstein College of Medicine, 1300 Morris Park Avenue, Rose F. Kennedy Center, Room 825, Bronx, NY 10461. **Author Contributions:** AUC, CFC, IVD, and BAJ designed and performed postsynaptic fractionation in autism mouse models. HEB and TAN performed tandem-mass-tag mass spectrometry and protein identification. AUC and IVD performed bioinformatics analysis and Western blots. ACA and DTP developed and characterized Pten haploinsufficiency mouse model. RAR developed and characterized Cacna1c G406R mouse model. AUC, CFC, and BAJ interpreted results and wrote the paper.

**Keywords:** proteomic, PSD, autism, mouse model, Rho GTPase, Rac, tandem mass tags, TMT

## Abstract

Impaired synaptic function is a common phenotype in animal models for autism spectrum disorder (ASD), and ASD risk genes are enriched for synaptic function. Here we leverage the availability of multiple ASD mouse models exhibiting synaptic deficits and behavioral correlates of ASD and use quantitative mass spectrometry with isobaric tandem mass tagging (TMT) to compare the hippocampal synaptic proteomes from 7 mouse models. We identified common altered cellular and molecular pathways at the synapse, including changes in Rho family small GTPase signaling, suggesting that it may be a point of convergence in ASD. Comparative analyses also revealed clusters of synaptic profiles, with similarities observed among models for Fragile X syndrome (*Fmr1* knockout), PTEN hamartoma tumor syndrome (*Pten* haploinsufficiency), and the BTBR+ model of idiopathic ASD. Opposing changes were found in models for cortical dysplasia focal epilepsy syndrome (*Cntnap2* knockout), Phelan McDermid syndrome (*Shank3* InsG3680), Timothy syndrome (*Cacna1c* G406R), and ANKS1B syndrome (*Anks1b* haploinsufficiency), which were similar to each other. We propose that these clusters of synaptic profiles form the basis for molecular subtypes that explain genetic heterogeneity in ASD despite a common clinical diagnosis. Drawn from an internally controlled survey of the synaptic proteome across animal models, our findings support the notion that synaptic dysfunction in the hippocampus is a shared mechanism of disease in ASD, and that Rho GTPase signaling may be an important pathway leading to disease phenotypes in autism.

## 1 Introduction

Autism spectrum disorder (ASD) is a neurodevelopmental disorder characterized by social-communication deficits and restrictive and repetitive behaviors. Although the specific cause of ASD is unknown, autism is highly heritable, with a monozygotic twin concordance rate of 40-80% (Gaugler et al. 2014). The genetic architecture of ASD is extraordinarily complex, with common inherited variants and rare *de novo* mutations working together to confer genetic risk (Weiner et al. 2017). This polygenic etiology presents challenges for elucidating the molecular pathogenesis of autism. However, ASD-related syndromes with defined genetic causes for autistic phenotypes present the best opportunities for elucidating the underlying mechanisms of ASD and identifying possible therapeutic targets (Sztainberg and Zoghbi 2016). These monogenic syndromes include Fragile X syndrome (*FMR1*), Rett syndrome (*MECP2*), PTEN hamartoma tumor syndrome (*PTEN*), tuberous sclerosis complex (*TSC1*/*TSC2*), Phelan McDermid syndrome (*SHANK3*), cortical dysplasia focal epilepsy syndrome (*CNTNAP2*) and Timothy syndrome (*CACNA1C*). Animal models of these syndromes have proven essential for studying the underlying neuropathology of ASD, especially changes in the complex processes of mammalian brain development and function.

Studies in rodents have shown that ASD risk genes converge on transcription regulation, protein homeostasis, and synaptic structure and function (Ruzzo et al. 2019; Pinto et al. 2014; De Rubeis et al. 2014). Accordingly, mice demonstrating the loss of *Fmr1, Mecp2, Pten, Tsc1/Tsc2, Shank* genes, or the *Nrxn* and *Nlgn* families all demonstrate changes in synaptic excitability or plasticity, and most also show altered dendritic growth or spine dynamics (Verma et al. 2019; Varghese et al. 2017; Hulbert and Jiang 2016; Huang, Chen, and Page 2016). Non-syndromic ASD models, such as those induced by valproic acid or maternal immune activation, also reveal structural and functional synaptic deficits, showing that environmental factors can lead to similar synaptic phenotypes (Andoh et al. 2019; Cellot et al. 2016; Li et al. 2018; Martin and Manzoni 2014; Patrich et al. 2016; Sui and Chen 2012; Wang et al. 2018). We recently developed a mouse model for ANKS1B haploinsufficiency, a rare genetic syndrome of ASD and other neurodevelopmental disorders caused by loss of the *ANKS1B* gene (Carbonell et al. 2019). *ANKS1B* was previously identified in ASD risk gene networks (Li et al. 2014), and we found that its product AIDA-1 regulates activity-induced protein synthesis (Jordan et al. 2007), hippocampal synaptic plasticity (Tindi et al. 2015), and NMDA receptor subunit composition (Tindi, et al. 2015; Carbonell, et al. 2019). AIDA-1 is a core protein of the postsynaptic density and interacts with PSD95 in a complex that contains other factors associated with neurodevelopmental disorders, including *Grin2b, Syngap1*, and *Nlgn*s (Carbonell, et al. 2019; Kaizuka and Takumi 2018). These findings support the idea that molecular mechanisms regulating synaptic function underlie ASD pathobiology (Bourgeron 2015; Zoghbi and Bear 2012).

Despite the heterogenous genetic architecture of autism and a complex etiology with contributions from environmental factors, ASD is diagnosed by distinct clinical criteria. Therefore, convergent cellular processes at the circuit, synaptic, or molecular level could underlie these shared behavioral phenotypes (Sestan and State 2018). In seeking convergent mechanisms among syndromic and non-syndromic forms of autism, comparative studies often narrowly focus on selected behaviors, specific synaptic phenomenology, or shared responses to preclinical pharmacological interventions (Schoen et al. 2019; Heise et al. 2018; Barnes et al. 2015). While broader comparisons have been made using transcriptomic analyses (Hernandez et al. 2020; Fores-Martos et al. 2019; Quesnel-Vallieres et al. 2019), these changes may not reflect mechanisms of disease due to multiple downstream levels of regulation such as protein translation, degradation, and transport (Vogel and Marcotte 2012). Proteomic approaches can therefore yield dramatically different results from transcriptomic profiles, as in a mouse model of Rett syndrome (Pacheco et al. 2017), and is particularly important when risk factors primarily regulate protein translation (*Fmr1, Tsc1*/*Tsc2, Pten*) or degradation (*Ube3a*) (Louros and Osterweil 2016). ASD models in which synaptic scaffolding and membrane localization are altered (*Shank3*, BTBR+) can also be investigated at the proteomic level, especially when the synaptic compartments and complexes are isolated by fractionation or immunoprecipitation (Reim et al. 2017; Lee et al. 2017; Murtaza, Uy, and Singh 2020; Brown et al. 2018). Indeed, analysis of postmortem brain tissue from ASD patients showed that some transcriptomic changes were not observed in cortical or cerebellar proteomes. However, changes in synaptic proteins predicted by ASD risk genes and animal models were confirmed (Abraham et al. 2019).

Here we compare the postsynaptic proteomes of seven mouse models for autism in which synaptic deficits have been described, with an emphasis on models of ASD-related syndromes: Fragile X syndrome (*Fmr1* −/Y), PTEN hamartoma tumor syndrome (*Pten* +/−), cortical dysplasia focal epilepsy syndrome (*Cntnap2* −/−), Phelan-McDermid syndrome (*Shank3* InsG3680), Timothy syndrome (*Cacna1c* G406R), ANKS1B syndrome (*Anks1b* +/−), and the BTBR+ inbred strain. These models have demonstrated face validity for autism, displaying hallmark behavioral correlates of ASD including social interaction deficits and restrictive behaviors (Kazdoba, Leach, and Crawley 2016; Zhou et al. 2016; Kabitzke et al. 2018; Carbonell, et al. 2019). In each model, we find evidence for upstream regulators that alter synaptic composition with predicted functional effects consistent with clinical phenotypes in ASD and individual syndromes. Mouse models can be clustered according to similarities and differences in upstream regulators, functional effects, and canonical pathways predicted by changes in their synaptic proteomes. Notably, we identified two groups of models with shared changes in the synaptic proteome, which may represent molecular subtypes of ASD. Proteins involved in Rho family GTPase signaling are commonly altered, especially the Rac signaling pathway. Measuring the expression of Rho GTPases in these models yielded new insights into differential regulation of Rho GTPase activity at the synapse. Appreciation of convergent synaptic changes underlying ASD is critical for defining pathogenic mechanisms and prioritizing treatments that can have broad efficacy (Sestan and State 2018). Our results suggest that targeting Rho GTPases may lead to the design of effective therapeutic interventions for diverse forms of autism spectrum disorder.

## 2 Materials and Methods

### 2.1 Mouse tissue and fractionation

Animals from the *Fmr1* knockout (stock #003025, MGI:1857169), *Cntnap2* knockout (stock #017482, MGI: 2677631), and *Shank3* InsG3680 mutant (stock #028778, MGI: 5775620) mouse models; and the BTBR+ (stock #002282), B6129SF2/J (stock #101045), and C57BL/6J (stock #000664) mouse strains were purchased from the Jackson Laboratory. Mice with *Pten* haploinsufficiency (MGI:2151804) and wild-type mice on the C57BL/6J background were obtained from the Page lab at the Scripps Research Institute Florida. Mice with the *Cacna1c* G406R mutation (MGI:5296904) and wild-type mice on the C57BL/6J background were obtained from the Rasmusson lab at the University at Buffalo. Heterozygotes from the *Anks1b* conditional knockout line (stock #035048, MGI:5779292) (Tindi, et al. 2015) and wild-type mice from the *Nestin-cre* transgenic line (stock #003771, MGI:2176173) were bred in house (Carbonell, et al. 2019). Mice from this line expressing the *Nestin-cre* transgene were genotyped for the *Anks1b^wt^* allele (forward: 5’-CACCCACAGCTCCATAGACAG-3’, reverse: 5’-GCACCTATTCCCTTCACCCTG-3’) and *Anks1b^fl^* allele (forward: 5’-AGTTGCCAGCCATCTGTTGT-3’, reverse: 5’-GGGTTCCGGATCAGCTTGAT-3’). For each proteomics experiment, we pooled hippocampi from 2 male mice of each genotype at 6-10 weeks of age. Postsynaptic density (PSD) enriched fractions were isolated from hippocampal tissue using sucrose density centrifugation and detergent extraction. To obtain sufficient material from hippocampal PSD fractionation of 2 animals, we used a modified protocol with a single incubation in 0.5% Triton-X100 as previously done (Tindi, et al. 2015).

### 2.2 Western blot

SDS-PAGE and Western blot were performed under standard conditions using the LI-COR system with the following antibodies: rabbit anti-Rac1 1:1000 (Proteintech #24702-1-AP), rabbit anti-RhoA 1:1000 (ABclonal), rabbit anti-Cdc42 (ABclonal), anti-Arf1 1:1000 (clone ARFS 1A9/5, Santa Cruz #53168), rabbit anti-Arf1 1:1000 (Proteintech #20226-1-AP), mouse anti-PSD95 1:1000 (NeuroMab), and rat anti-tubulin 1:1000 (Cell Signaling Tech). Statistical analysis for Western blot was performed in JMP 14 (SAS) as in Supplementary Table 6**Error! Reference source not found.**.

### 2.3 Tandem mass tag labeling of peptides

For each experiment, tryptic peptide pools generated from 10 hippocampal PSD-enriched samples (15 μg protein each for Experiment 1, 20 μg protein each for Experiment 2) were reacted with unique isobaric labels within a tandem mass tags (TMT) set (Thompson et al. 2003) and analyzed together using high-resolution (Q Exactive HF) mass spectrometry (Carbonell, et al. 2019; Klein et al. 2019). Each sample was electrophoresed briefly (dye front 5 mm) into a 4-12% SDS-PAGE gel. The gel was washed 3x in ddH_2_O for 15 min each and visualized by staining overnight with GelCode® Coomassie blue reagent (Pierce). Stacked protein bands were excised from the gel, reduced with DTT, and alkylated with iodoacetamide. In-gel digestion was performed using 5 ng/μL mass spectrometry-grade trypsin (Trypsin Gold, Promega) in 50 mM NH_4_HCO_3_ digestion buffer. The resulting peptides were desalted using a Stage Tip manually packed with Empora C18 High Performance Extraction Disks (3M) (Rappsilber, Mann, and Ishihama 2007) and eluted peptide solutions were dried under vacuum.

Peptides were then resuspended in 18 μL acetonitrile (ACN), and 57 μL of 0.2 M HEPES pH 8.5 was added to each sample. TMT10-plex amine reactive reagents (Thermo Fisher, 5 mg per vial) were re-suspended in 1024 μL anhydrous acetonitrile and 25 μL of reagent was added to each sample (TMT label: peptide [w/w] =12:1) and mixed briefly by vortexing. The mixture was incubated at RT for 1 hr, quenched by the addition of 10 μL 5% hydroxylamine for 15 min, and acidified by the addition of 10 μL 10% formic acid. A 5-μL aliquot from each reaction was desalted on a StageTip, analyzed by LC-MS/MS with a Q Exactive Orbitrap HF (high field), and the resulting spectra searched with MaxQuant using its corresponding TMT label as variable modifications on N-terminus and lysine. The percentage of peptides with either N-terminal or lysine TMT labels was calculated, indicating the labeling efficiency for each channel. Labeling efficiency was 95% or greater for each channel. To ensure that equal amounts of labeled peptides from each channel were mixed together, a two-step mixing strategy was employed: in the first step, an identical ∼1 μL volume of peptides from each channel was mixed and analyzed, and the value of the median ratio (median of the ratios of all peptide intensities of one channel over their corresponding peptide average intensities of all channels) for each channel was determined as the correction factor. In the second step, the rest of the peptides were mixed by adjusting their volume using the correction factors. In this way, median ratios ranging from 0.97 to 1.02 were achieved as previously reported (Erdjument-Bromage, Huang, and Neubert 2018). The final mixture of reaction products from 10 TMT channels were desalted on a Sep-Pak tC18 1mL Vac Cartridge (Waters, #WAT03820). Eluted peptides were dried by vacuum centrifugation and stored at −20°C.

### 2.4 Hydrophilic Interaction Liquid Chromatography (HILIC) Fractionation of Peptides

We fractionated PSD samples by offline HILIC to increase depth of coverage and decrease ratio compression due to co-fragmenting peptides (Huang et al. 2017). Final TMT mix was dissolved in 90% acetonitrile with 0.1% TFA and peptide separation was carried out on an Agilent pump equipped with a TSK gel amide-80 column (4.6mm ID, 25cm long) from TOSOH Bioscience, LLC, PA, USA). A gradient of 90% acetonitrile with 0.1% TFA was introduced over 65 min., and a fraction was collected every 2 minutes. Concatenated pools of peptides (10 pools) were finally created by pooling non-adjacent peptide fractions; about 10% of each pool was used for LC-MS/MS analysis.

### 2.5 Liquid chromatography-tandem mass spectrometry

Online chromatography was performed with a Thermo Easy nLC 1000 ultrahigh-pressure UPLC system (Thermo Fisher) coupled online to a Q Exactive HF with a NanoFlex source (Thermo Fisher). Analytical columns (∼23 cm long and 75 μm inner diameter) were packed in house with ReproSil-Pur C18 AQ 3 μM reversed-phase resin (Dr. Maisch GmbH, Ammerbuch-Entringen). The analytical column was placed in a column heater (Sonation GmbH, Biberach) regulated to a temperature of 45°C. The TMT peptide mixture was loaded onto the analytical column with buffer A (0.1% formic acid) at a maximum back-pressure of 300 bar. Peptides were eluted with a 2-step gradient of 3% to 40% buffer B (100% ACN and 0.1% formic acid) in 180 min and 40% to 90% B in 20 min, at a flow rate of 250 nL/min over 200 min using a 1D online LC-MS2 data-dependent analysis (DDA) method as follows: MS data were acquired using a data-dependent top-10 method, dynamically choosing the most abundant not-yet-sequenced precursor ions from the survey scans (300–1750 Th). Peptide fragmentation was performed via higher energy collisional dissociation with a target value of 1 × 10^5^ ions determined with predictive automatic gain control. Isolation of precursors was performed with a window of 1 Th. Survey scans were acquired at a resolution of 120,000 at *m*/*z* 200. Resolution for HCD spectra was set to 60,000 at *m*/*z* 200 with a maximum ion injection time of 128 ms. The normalized collision energy was 35. The underfill ratio specifying the minimum percentage of the target ion value likely to be reached at the maximum fill time was defined as 0.1%. Precursor ions with single, unassigned, or seven and higher charge states were excluded from fragmentation selection. Dynamic exclusion time was set at 30 sec. Each of the TMT 10-plex samples was analyzed in triplicate.

All data were analyzed with the MaxQuant proteomics data analysis workflow (version 1.5.5.7) with the Andromeda search engine (Cox et al. 2011; Tyanova, Temu, and Cox 2016). The type of the group specific analysis was set to Reporter ion MS2 with 10plex TMT as isobaric labels for Q Exactive HF MS2 data. Reporter ion mass tolerance was set to 0.01 Da, with activated Precursor Intensity Fraction (PIF) value set at 0.75. False discovery rate was set to 1% for protein, peptide spectrum match, and site decoy fraction levels. Peptides were required to have a minimum length of eight amino acids and a maximum mass of 4,600 Da. MaxQuant was used to score fragmentation scans for identification based on a search with an allowed mass deviation of the precursor ion of up to 4.5 ppm after time-dependent mass calibration. The allowed fragment mass deviation was 20 ppm. MS2 spectra were used by Andromeda within MaxQuant to search the Uniprot mouse database (01092015; 16,699 entries) combined with 262 common contaminants. Enzyme specificity was set as C-terminal to arginine and lysine, and a maximum of two missed cleavages were allowed. Carbamidomethylation of cysteine was set as a fixed modification and N-terminal protein acetylation, deamidated (N, Q) and oxidation (M) as variable modifications. The reporter ion intensities were defined as intensities multiplied by injection time (to obtain the total signal) for each isobaric labeling channel summed over all MS/MS spectra matching to the protein group as previously validated (Tyanova, Temu, and Cox 2016). Following MaxQuant analysis, the protein and peptide .txt files were imported into Perseus (version 1.5.6.0) software which was used for statistical analysis of all the proteins identified.

### 2.5 Bioinformatics analysis

For each mouse model of autism, fold-change of each protein was calculated by dividing relative abundance of each sample by the abundance in the appropriate control sample. For the *Fmr1, Pten, Cntnap2, Cacna1c*, and BTBR+ models, we used the average of 2 samples of C57BL/6J mice as the control value. For *Shank3* mice on the B6129SF2/J background, we used a sample of wild-type mice from the same colony as the control value. For the *Anks1b* Het mice on the *Nestin-cre* C57BL/6J background, we used wild-type littermates on the *Nestin-cre* C57BL/6J background as the control value. Of the proteins quantified in all samples, we filtered duplicate genes for the protein IDs with highest number of unique peptides, highest sequence coverage, and lowest Q-values, yielding the highest intensity and scores. We considered the proteins identified by >2 peptides in Experiment 1 or >9 peptides in Experiment 2 as the hippocampal PSD proteome, to obtain a reliable fold change for each protein. We used this list of 788 proteins (Experiment 1) or 830 proteins (Experiment 2) as input for multiple proteins in StringDB under the species *Mus musculus*. Raw data for gene ontology (GO) enrichments were generated in StringDB (Szklarczyk et al. 2015).

Functional annotation, activation prediction, and regulatory network construction was performed for each mouse model using Ingenuity Pathway Analysis (IPA, QIAGEN Bioinformatics, version 52912811). Fold-change values for each protein in the PSD proteome was used as input for a *Core Analysis* in IPA using the following default settings: Expression Analysis; General Settings = Ingenuity Knowledge Base (Genes Only), Direct Relationships; Networks = Interaction networks, Include endogenous chemicals; Node Types = All; Data Sources = All; Confidence = Experimentally Observed; Species = All; Tissues & Cell Lines = All; Mutations = All. For each model, Regulator Effects networks were generated to yield regulators of Type = All, diseases and functions of Category = All, with a *p*-value cutoff of <0.01 and *z*-score cutoff of >|2|. To score similarity between models overall and in each domain (Upstream Regulators, Downstream Effects, and Canonical Pathways), *Analysis Match* was used. In *Analysis Match*, activity signatures for each *Core Analysis* are generated by taking the top 50 (Upstream Regulators and Downstream Effects) or 10 (Canonical Pathways) entities that are activated (*z*-score >2) and inhibited (*z*-score <-2) in each domain. Similarity *z*-scores between Core Analyses are defined using the following formula, where *N* is the total number of overlapping entities in each analysis, *N*_+_ is the number of correct matches, and *N*_−_the number sof incorrect matches:

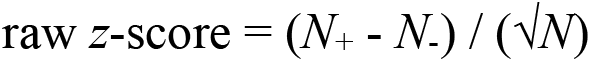

The raw *z*-score is divided by a hypothetical perfect match (*N*_+_ = *N* ≤ 100, *N*_−_= 0) and multiplied by 100% to obtain a normalized *z*-score, where a score of 100 is a perfect match to itself. Raw and normalized *z*-scores are negative when *N*_−_> *N*_+_, yielding an opposite activation signature. To generate overall *p*-value scores, the −log_10_ of the *p*-values (maximum value of 50) were calculated for each domain, and expressed as a percentage of the maximum possible −log_10_*p* (a perfect match to itself). To compare models, the *Core Analysis* of each model were used as input for a *Comparison Analysis* in IPA. Hierarchical analysis was used to cluster annotations in Upstream Analysis, Downstream Effects, and Canonical Pathways, and to cluster models for similarity relationships.

## 3 Results

### 3.1 TMT-MS analyses of postsynaptic density fractions from mouse models of autism

To investigate the synaptic proteome in autism, we selected 7 mouse models: *Fmr1* knockout (Fmr1 KO) (Consortium 1994), *Pten* haploinsufficiency (Pten Het) (Clipperton-Allen and Page 2015; Clipperton-Allen and Page 2014), *Cntnap2* knockout (Cntnap2 KO) (Penagarikano et al. 2011), *Shank3* frameshift mutation InsG3680 (Shank3*) (Zhou, et al. 2016), Timothy syndrome mutation *Cacna1c* G406R (Cacna1c*) (Bader et al. 2011), *Anks1b* haploinsufficiency (Anks1b Het) (Carbonell, et al. 2019), and the BTBR+ inbred strain (McFarlane et al. 2008). All selected models display abnormal behaviors in domains relevant to ASD, including social approach and interaction, stereotyped movements, learning and memory, and sensorimotor function. These models have also been used to illustrate synaptic dysfunction, including altered synapse formation, excitatory/inhibitory balance, synaptic plasticity, and glutamatergic signaling (Hulbert and Jiang 2016; Varghese, et al. 2017; Bagni and Zukin 2019; Carbonell, et al. 2019). To analyze changes in the synaptic proteome for each mouse model of autism, we isolated the hippocampus and used sucrose density-based centrifugation to yield postsynaptic density (PSD) enriched fractions. Western blot of these samples demonstrated enrichment for the marker PSD95 in each PSD fraction compared to total hippocampal lysate from each model (Figure 1**A**). We then processed the samples for tandem-mass-tag mass spectrometry (TMT-MS) to simultaneously analyze the synaptic proteome in each ASD model with other models and their appropriate control samples (Figure 1**B**). This method reduces the variability induced by mass spectrometry and allows comparison of protein abundance from peptides reliably quantified with a minimum of 2 peptides, across all samples (**Error! Reference source not found.**). After peptide identification and filtering for proteins by number of peptides quantified, we obtained a synaptic proteome of 788 proteins for Experiment 1 (Supplementary Table 1) and 830 proteins for Experiment 2 (Supplementary Table 2). This number is consistent with previous studies showing 984 PSD proteins in mice, 748 in humans, and 546 consensus proteins between them (Bayes et al. 2012). As expected, gene ontology (GO) analysis of the synaptic proteomes we identified in STRING (Szklarczyk et al. 2019) revealed significant enrichment for synaptic proteins and similar enrichments between experiments (Figure 1**C**). As a functional readout, the synaptic proteomes were also enriched for reaction pathways involved in synaptic formation and transmission (Figure 1**D**Error! Reference source not found.). Consistent with the organized structure and specialized function of the PSD, biological processes related to cellular organization and neural development were enriched (Supplementary Table 3). Comparing the synaptic proteomes to the 1,104 synaptic proteins annotated in the SynGO database (Koopmans et al. 2019), 336 synaptic proteins were identified in Experiment 1 (Fisher’s exact test, *p* = 2.09e-188) and 295 synaptic proteins were found in Experiment 2 (Fisher’s exact test, *p* = 5.23e-142) (SynGO geneset analysis, https://syngoportal.org). These results validate the efficient detection of synaptic proteins through TMT-MS.

**Table 1.** Pathways related to Rho GTPase signaling are differentially activated in the synapse across mouse models of autism. Results from six Canonical Pathways in IPA related to Rho GTPase signaling are shown for ASD mouse models in Experiments 1 and 2. Overlap −log_10_(*p*-values) are based on the enrichment of pathway components observed in each proteome from Fisher’s exact test, and ratio is the fraction of pathway components observed: these values are constant within each experiment. Activation *z*-scores are predicted based on observed protein fold changes in each model and differ based on altered proteome composition in each model and experiment.

| Canonical Pathway | EXPERIMENT 1 |  |  | EXPERIMENT 2 |  |  |
| --- | --- | --- | --- | --- | --- | --- |
| | $-\log_{10}p$ (ratio) | Model | z-score | $-\log_{10}p$ (ratio) | Model | z-score |
| Signaling by Rho Family GTPases | 17.6 (0.172) | Fmr1 KO | 4.70 | 18.9 (0.184) | Fmr1 KO | 0.82 |
|  |  | Pten Het | 3.31 |  | Pten Het | 5.43 |
|  |  | BTBR+ | 0.17 |  | BTBR+ | 2.80 |
|  |  | Cntnap2 KO | -0.17 |  | Cntnap2 KO | 0.82 |
|  |  | Shank3* | 0.52 |  | Anks1b Het 1 | -2.47 |
|  |  | Cacna1c* | -4.35 |  | Anks1b Het 2 | -4.77 |
| Rac Signaling | 12.5 (0.214) | Fmr1 KO | 3.96 | 8.5 (0.179) | Fmr1 KO | 2.98 |
|  |  | Pten Het | 3.13 |  | Pten Het | 3.44 |
|  |  | BTBR+ | 1.04 |  | BTBR+ | 2.52 |
|  |  | Cntnap2 KO | 0.21 |  | Cntnap2 KO | 0.69 |
|  |  | Shank3* | 0.63 |  | Anks1b Het 1 | -0.69 |
|  |  | Cacna1c* | -3.96 |  | Anks1b Het 2 | -2.98 |
| RhoA Signaling | 13.3 (0.211) | Fmr1 KO | 3.00 | 19.3 (0.268) | Fmr1 KO | 1.26 |
|  |  | Pten Het | 3.40 |  | Pten Het | 3.41 |
|  |  | BTBR+ | 1.80 |  | BTBR+ | 4.13 |
|  |  | Cntnap2 KO | 1.40 |  | Cntnap2 KO | 1.98 |
|  |  | Shank3* | 1.80 |  | Anks1b Het 1 | -2.34 |
|  |  | Cacna1c* | -2.60 |  | Anks1b Het 2 | -2.69 |
| Cdc42 Signaling | 4.85 (0.108) | Fmr1 KO | 2.36 | 5.57 (0.120) | Fmr1 KO | 3.00 |
|  |  | Pten Het | 3.77 |  | Pten Het | 1.50 |
|  |  | BTBR+ | 2.36 |  | BTBR+ | 1.50 |
|  |  | Cntnap2 KO | 2.36 |  | Cntnap2 KO | 1.00 |
|  |  | Shank3* | 0.94 |  | Anks1b Het 1 | -0.50 |
|  |  | Cacna1c* | -2.36 |  | Anks1b Het 2 | -1.50 |
| RhoGDI Signaling | 17.4 (0.200) | Fmr1 KO | -4.91 | 13.9 (0.183) | Fmr1 KO | -1.80 |
|  |  | Pten Het | -3.40 |  | Pten Het | -3.80 |
|  |  | BTBR+ | -0.38 |  | BTBR+ | -2.20 |
|  |  | Cntnap2 KO | -0.38 |  | Cntnap2 KO | -1.00 |
|  |  | Shank3* | -0.38 |  | Anks1b Het 1 | 1.00 |
|  |  | Cacna1c* | 3.78 |  | Anks1b Het 2 | 3.40 |
| Regulation of Actin-based Motility by Rho | 11.4 (0.223) | Fmr1 KO | 2.98 | 6.63 (0.170) | Fmr1 KO | 2.32 |
|  |  | Pten Het | 3.44 |  | Pten Het | 1.81 |
|  |  | BTBR+ | 2.52 |  | BTBR+ | 2.84 |
|  |  | Cntnap2 KO | 1.61 |  | Cntnap2 KO | 2.32 |
|  |  | Shank3* | 1.15 |  | Anks1b Het 1 | -1.29 |
|  |  | Cacna1c* | -2.07 |  | Anks1b Het 2 | -1.81 |

**Figure 1.**
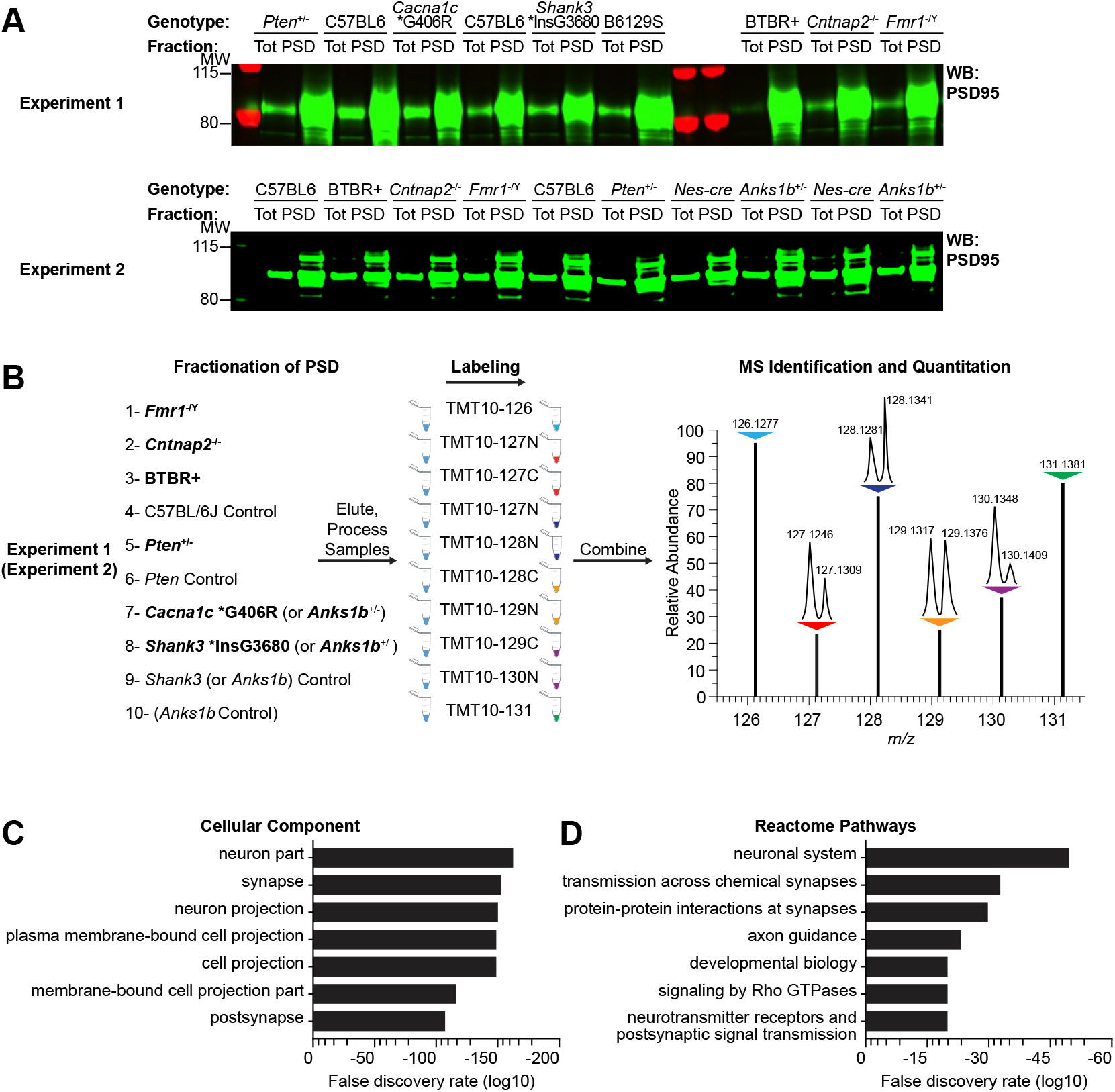
Fractionation of mouse hippocampus yields the postsynaptic proteome. **A**) Total lysate and postsynaptic density (PSD) enriched fractions from mouse models of autism spectrum disorder (ASD) demonstrate qualitative enrichment for the synaptic marker PSD95 by Western blot (4 μg each sample). **B**) Comparative proteomics scheme showing parallel processing of hippocampal synaptic fractions from ASD mouse models and their control samples. Shank3* and Cacna1c* models were only assayed in Experiment 1, and both Anks1b Het samples were assayed in Experiment 2. **C**) Gene ontology (GO) analysis of the postsynaptic proteome in STRING (Experiment 1) showed enrichment for cellular components and reactome pathways (**D**) expected for synaptic fractions.

### 3.2 Predicted effects drive synaptic similarities among ASD mouse models

To obtain a fold-change of the synaptic proteins for each ASD model, we used C57BL/6J mice as controls, except for the Shank3* and Anks1b Het models, for which we used wild-type animals from the same colony on the B6129SF2/J and *Nestin-cre* C57BL/6J backgrounds, respectively. For each mouse model, we performed a *Core Analysis* of fold-changes in the synaptic proteome in Ingenuity Pathway Analysis (IPA, QIAGEN Bioinformatics) to score the Diseases and Functions (Supplementary Table 4), Upstream Regulators (Supplementary Table 5), and Canonical Pathways (Supplementary Table 5) significantly enriched (*p*-values) and altered (*z*-scores) in the synaptic proteomes from Experiment 1 and Experiment 2 (Kramer et al. 2014). To test whether ASD mouse models display shared changes in their synaptic proteomes, we quantitatively compared the results of the *Core Analysis* from each model using *Analysis Match* in IPA (Figure 2**A-B**). To obtain overall similarity scores, IPA incorporates activation *z*-scores from Upstream Regulators, Downstream Effects (Diseases and Functions), and Canonical Pathways from each *Core Analysis* into activation patterns (activity signatures) that can be compared across analyses (see **Materials and Methods**). Similarity *z*-scores reflect the percentage of maximum similarity to a given model (out of 100%, a perfect match to itself).

**Figure 2.**
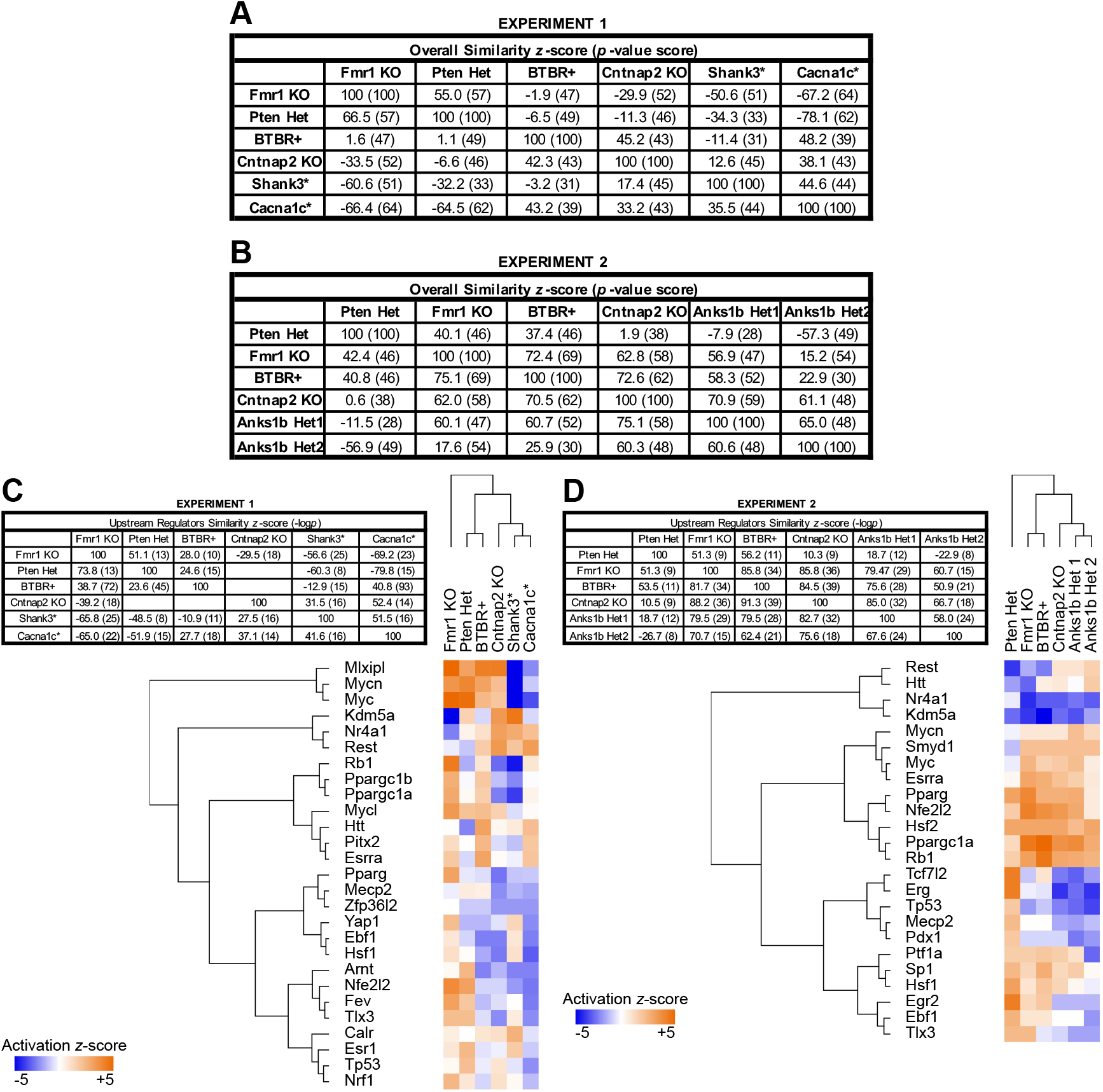
Overall similarities and differences among mouse models of ASD reflect patterns in predicted Upstream Regulators. **A**) Analysis Match in IPA scores the similarities in activation patterns among Upstream Regulators, Downstream Effects, and Canonical Pathways between ASD mouse models in Experiments 1 and 2. (**B**) Overall similarity scores in Experiment 1 and Experiment 2 show congruent relationships among the Fmr1 KO, Pten Het, BTBR+, and Cntnap2 KO models examined in both experiments, and for both samples of the Anks1b Het model in Experiment 2. Cacna1c* and Shank3* mutant models were only assessed in Experiment 1. **C**) For Upstream Regulators alone, the same pattern of similarity among Fmr1, Pten Het, and BTBR+; as opposed to Cntnap2 KO, Shank3*, Cacna1c*, and Anks1b Het, is observed. Clustering of ASD mouse models shows the relationships among models in Experiment 1 and (**D**) Experiment 2. Tree diagrams show hierarchical cluster analysis of Upstream Regulators in IPA with activation *z*-scores >|2| and enrichment −log_10_(*p*-values) >2 (Benjamin-Hochberg corrected for multiple comparisons).

Overall similarity *z*-scores and *p*-value scores (out of 100) from Experiment 1, where we tested 6 ASD models, are given in Figure 2**A**. For the Fmr1 KO mouse model, the highest similarity *z*-score was with Pten Het mice (*z*-score 55.0). The Cacna1c* (*z*-score −67.2) and Shank3* (*z*-score −50.6) were the most dissimilar models to the Fmr1 KO, with significant changes in the opposite direction to those observed in the Fmr1 KO mouse indicated by high negative *z*-scores. Similarly, the Pten Het model was most similar to Fmr1 KO, and most dissimilar to the Cacna1c* and Shank3* mice. For the BTBR+ strain, the Cntnap2 KO (*z*-score 45.2) and Cacna1c* (*z*-score 48.2) were both similar and no other models were substantially dissimilar. For the Cntnap2 KO model, the BTBR+ strain (*z*-score 42.3) and Cacna1c* model (*z*-score 38.1) were most similar and the Fmr1 KO model most dissimilar (*z*-score −33.5). For the Shank3* model, the Cacna1c* model was most similar (*z*-score 44.6), with Fmr1 KO (*z*-score −60.6) and Pten Het (*z*-score −32.2) models showing opposite activation signatures. For the Cacna1c* model, activation patterns were strongly opposite in direction to Fmr1 KO (*z*-score −66.48) and Pten Het (*z*-score −64.5), while somewhat similar to other models (BTBR *z*-score 43.2, Shank3* *z*-score 35.5, Cntnap2 KO *z*-score 33.2). In the overall similarity *z*-scores, a pattern emerged of consistent agreement between Fmr1 KO and Pten Het models, intermediate similarity of BTBR+ and Cntnap2 KO mice to other models, and dissimilarity of Shank3* and Cacna1c* models to Fmr1 KO and Pten Het (Figure 2**A**).

To test for reproducibility of findings, we performed a second experiment (Experiment 2), where we once again compared the Fmr1 KO and BTBR+ strains, which are widely studied mouse models of monogenic and idiopathic ASD, to Pten Het and Cntnap2 KO mice, which showed the most similarity to these models in Experiment 1. This time, we included biological replicates of our Anks1b Het mouse model (Figure 2**B**). Similar to Experiment 1, the Pten Het model was most similar to Fmr1 (*z*-score 40.1), although Fmr1 KO was also similar to BTBR+ (*z*-score 72.4) and Cntnap2 KO (*z*-score 62.8) models. Compared to Experiment 1, overall similarity among the replicated models was greater in Experiment 2, with BTBR+ similar to Cntnap2 KO (*z*-score 72.6) and to other models (Fmr1 KO *z*-score 75.1, Pten Het *z*-score 40.8). Cntnap2 KO was also similar to BTBR+ (*z*-score 70.5) and Fmr1 KO (*z*-score 62.0), although not similar to Pten Het (*z*-score 0.6). Strikingly, the Anks1b Het samples were most consistently similar to Cntnap2 KO (Anks1b Het1 *z*-score 70.9, Anks1b Het2 *z*-score 61.1) and had opposite activation signatures to Pten Het (Anks1b Het1 *z*-score −7.9, Anks1b Het2 *z*-score −57.3). Similarities to Fmr1 KO and BTBR+ were not consistent but tended to be positive (Figure 2**B**). Biological replicates for the Anks1b Het model were highly similar to each other, demonstrating internal consistency and thus validating our TMT methodology and IPA-based predictions from proteomics measurements (Anks1b Het1 to Het2 *z*-score 65.0, Het2 to Het1 *z*-score 60.6). For all similarity *z*-scores, *p*-values for overlap of commonly activated and inhibited entities were calculated in IPA using Fisher’s exact test (*p*<0.01 for all comparisons).

The overall similarity *z*-scores incorporate activation patterns from predicted Upstream Regulators, Downstream Effects, and Canonical Pathways. For each domain of comparison, the trends for similar or opposite activation between ASD mouse models were largely consistent. When comparing Upstream Regulators, activation signatures in Fmr1 KO and Pten Het models were similar to each other and dissimilar to Shank3 and Cacna1c* by Analysis Match and cluster analysis (Figure 2**C**). While similar to Fmr1 KO and Pten Het in both Experiment 1 and 2 (Figure 2**C**), the BTBR+ strain also showed similarities to the Cacna1c* model (*z*-score 40.8). Experiment 2 showed similar activation patterns across models, except for the Pten Het model, which showed limited similarity to Cntnap2 KO (*z*-score 10.3) and Anks1b Het (Het1 *z*-score 18.7, Het2 *z*-score −22.9). In Experiment 1, predicted upstream regulators with the largest magnitude of activation or inhibition included Mlxipl, Myc, Kdm5a, and Rest. Variants in Mlxipl are associated with autism in Williams-Beuren syndrome (Codina-Sola et al. 2019). Activation *z*-scores for Mlxipl and Myc were driven by changes in ribosomal and mitochondrial proteins regulated by Myc and Ctnnb1 (Supplementary Table 5). Kdm5a is part of a family of histone demethylases, of which several members have been implicated in intellectual disability (Vallianatos et al. 2018; Zamurrad et al. 2018). While Kdm5a activation was inconsistent in Experiment 1 (Figure 2**C**), Experiment 2 showed Kdm5a inhibition across Pten Het, Fmr1 KO, BTBR+, Cntnap2 KO, and Anks1b Het models (Figure 2**D**). Across experiments, inhibition of Rest (RE1-silencing transcription factor) and activation of Tlx3 was predicted in Fmr1 KO and Pten Het, while the opposite was seen in Cntnap2 KO, Cacna1c*, and Anks1b Het models (Figure 2**C-D**). Mecp2 was predicted to be activated or unchanged in Fmr1 KO, Pten Het, and BTBR+, but inhibited in the other 4 models (Figure 2**C-D**).

### 3.3 Altered synaptic composition yields networks of functional effects

After comparing Upstream Regulators, we compared activation signatures for Downstream Effects on diseases and functions enriched in the synaptic proteomes. Like Upstream Regulators, Downstream Effects are predicted in IPA from an updated knowledge base drawn from the literature (Kramer, et al. 2014). Results were largely consistent with overall similarity trends, with significant similarities between Fmr1 KO and Pten Het by Analysis Match and cluster analysis (Figure 3**A-B**). However, there were fewer differences between BTBR+ and the other models than overall or in Upstream Regulators, and Shank3* showed activation patterns intermediate between the Fmr1 KO/Pten Het and BTBR+/Cntnap2/Cacna1c* clusters (Figure 3**A**). In Experiment 2, Anks1b Het mice showed similar activation signatures to Cntnap2 KO (Anks1b Het1 *z*-score 55.3, Anks1b Het2 *z*-score 57.1) and opposing activation to the Pten Het and Fmr1 KO models by Analysis Match and cluster analysis (Figure 3**B**). Across models, the most divergent functional effects were on cellular structure, including cellular protrusions, microtubule dynamics, and cytoskeleton regulation for neuronal development, which were activated for Fmr1 KO and Pten Het and inhibited for other models (Figure 3**A-B**). Opposing effects on disease were evident in movement and seizure disorders, which were predicted to be downregulated in Fmr1 KO and Pten Het mice (Figure 3**A-B**), while effects on cognitive impairment and emotional behavior did not strongly differ across models (Figure 3**A-B**).

**Figure 3.**
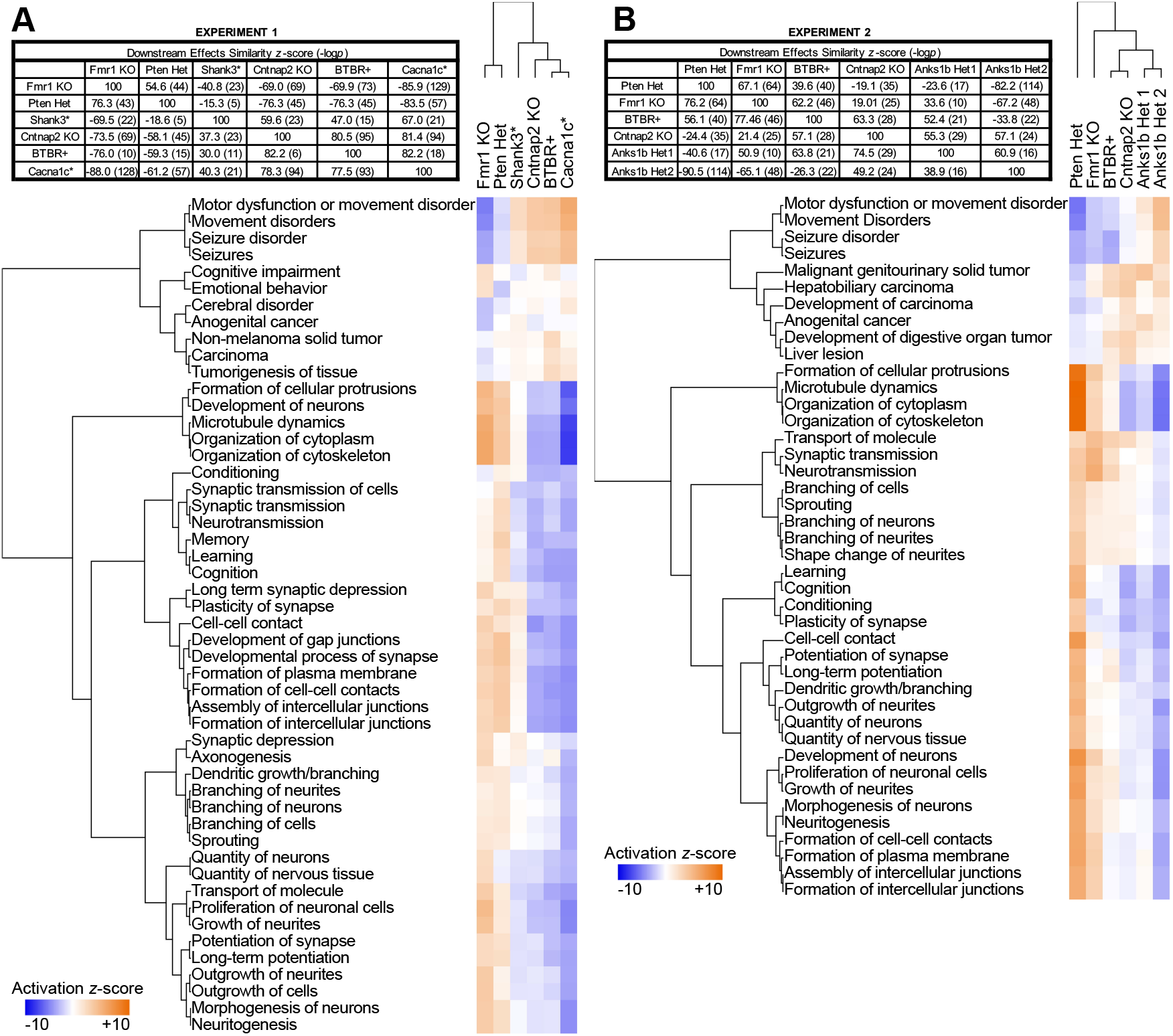
Predicted Downstream Effects on diseases and functions show patterns among mouse models that are consistent with Overall and Upstream Regulators. **A**) For Downstream Effects, similarity scores among ASD mouse models in Experiment 1 and Experiment 2 show similarities among the Fmr1 KO, Pten Het, BTBR+, and Shank3* model, as opposed to the Cntnap2 KO, Cacna1c*, and *Anks1b* Het model (**B**). Clustering of ASD mouse models shows the relationships among mouse models in Experiment 1 and Experiment 2. Tree diagrams show hierarchical cluster analysis of Downstream Effects in IPA with activation *z*-scores >|2| and enrichment −log_10_(*p*-values) >25 (Benjamini-Hochberg corrected for multiple comparisons).

We next focused on IPA results for Regulatory Effects, which links Upstream Regulators (genes, RNAs, and proteins, Figure 2**C-D**) to Downstream Effects (diseases and disorders, physiological systems, cellular and molecular functions, Figure 3**A-B**) through observed changes in the synaptic proteome. Top-scoring networks for each model display consistent directionality (score >0) and yield unified, functional representations of altered synaptic proteins in each model with implied upstream causes and predicted downstream effects from the IPA knowledge base. For the Fmr1 KO model, activated Tlx3, Erg, and Fev regulate synaptic targets that contribute to a range of neurodevelopmental functions, including synaptic development, motor coordination, cognitive impairment, and hyperactivity (Experiment 1, Figure 4**A**). A larger network driven by inhibited Kdm5a and Nr4a1, among others, predicts altered energy metabolism and congenital neurological disorder (Experiment 2, Figure 4**B**). These trends are consistent across experiments, with inhibited Kdm5a predicted in Experiment 1 (Error! Reference source not found.**A**) and activated Tlx3 in Experiment 2 (Error! Reference source not found.**B**). Similarly, the Pten Het model also showed predicted activation of Tlx3 and inhibition of Kdm5a, predicting increased neurotransmission and neuronal movement (Experiment 2, Figure 4**C**). Tlx3 activation of neurotransmission was also predicted as a top network in Experiment 1 (Error! Reference source not found.**C**). Inhibition of Rest and activation of Mecp2 were predicted to activate endocytosis and neurite outgrowth, as well as proteins involved in long-term depression of the synapse (Experiment 2, Figure 4**D**). Similar to the Pten Het model, the BTBR+ strain shows inhibition of Rest, predicting increased vesicle endocytosis and transport (Experiment 2, Figure 4**E**). Additionally, inhibition of Neurod1 promotes organismal death while reducing long-term potentiation and memory (Experiment 2, Figure 4**F**). However, BTBR+ strain diverges from the Fmr1 KO and Pten Het models in the predicted inhibition of Fev, leading to reduced neurotransmission downstream (Error! Reference source not found.**C**).

**Figure 4.**
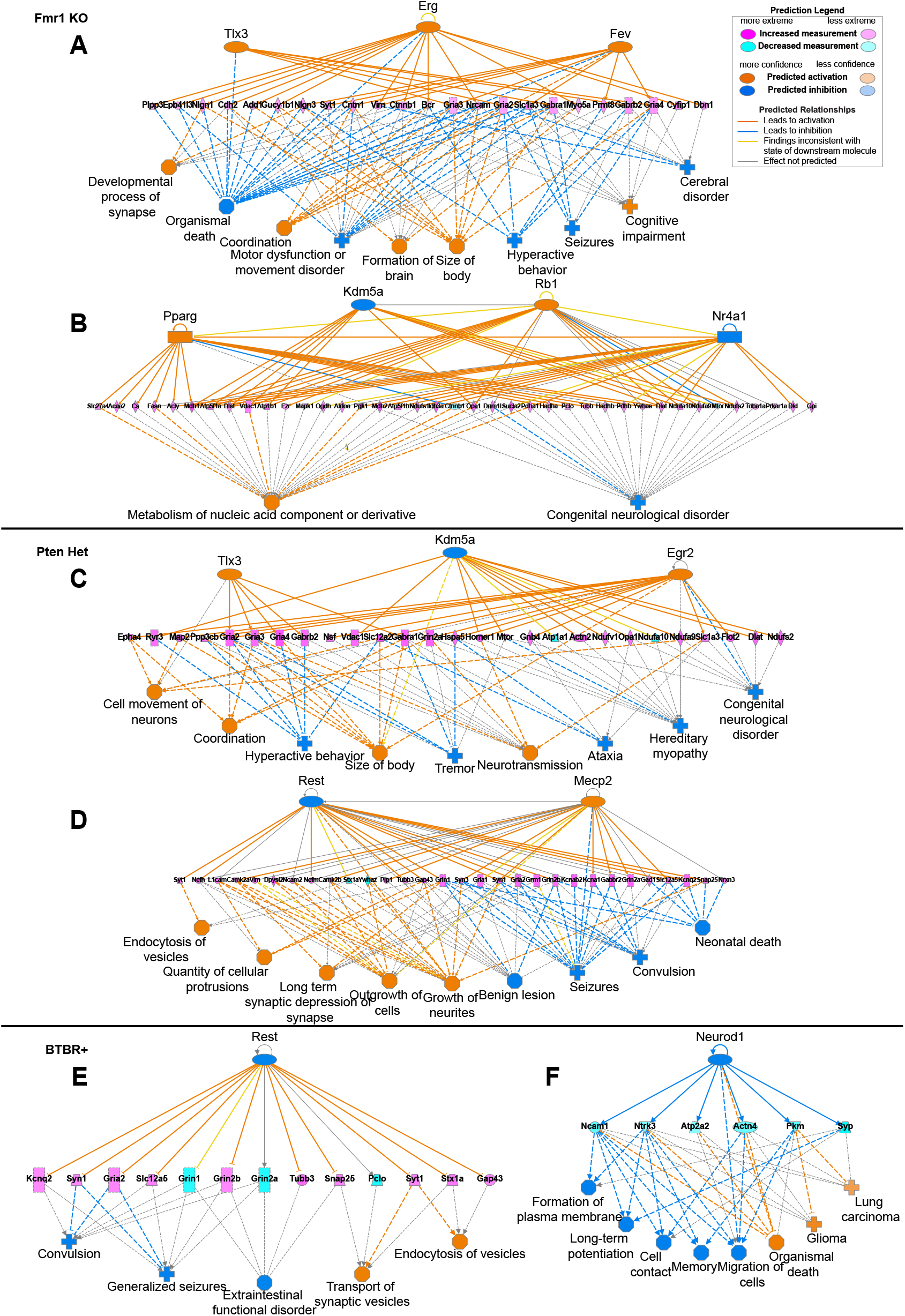
Altered synaptic composition predicts similar networks of functional effects in Fmr1 KO, Pten Het, and BTBR+ mice. **A-B**) Top regulatory networks for Fmr1 KO mice. **C-D**) Top regulatory networks for Pten Het mice. **E-F**) Top regulatory networks for BTBR+ mice. Upstream regulators and downstream effects were predicted in IPA from fold-changes of proteins in the postsynaptic proteome.

In contrast to Fmr1 KO and Pten Het, the Cntnap2 KO model showed inhibited expression of genes controlled by Tlx3 as the top network in both experiments, predicting reduced neurotransmission and hyperactive behavior (Figure 5**A,** Error! Reference source not found.**E**). Another regulatory network, driven by inhibited Mecp2 and Zpf36l2, was predicted to reduce neurite growth and vesicle quantity while promoting seizures (Experiment 1, Figure 5**B**). Downward changes of synaptic channels and receptors is consistent with activated Rest, which was predicted to reduce vesicle transport, synaptic plasticity, and cognition (Experiment 1, Figure 5**C**). In the Shank3* model, Zfp36l2 was also inhibited in a network with activated Calr, predicting neurodegeneration and altered protein metabolism (Figure 5**D**). Although changes in the Shank3* synapse were milder than in other models, inhibited Arnt and Tfe3 predicted motor dysfunction and movement disorders in another top network (Error! Reference source not found.**F**). In the Cacna1c* model, the top regulatory network shows downregulation of both Fev and Tlx3, predicting inhibition of neural development and synaptic transmission and promotion of seizures and motor dysfunction (Figure 5**E**). As in the BTBR+ strain, inhibition of Neurod1 predicts reduced synaptic plasticity and memory (Figure 5**F**). However, an extensive network driven by activated Rest and Htt and inhibited Nfe212 promotes heart failure and impaired synaptic depression, in contrast to the Fmr1, Pten Het, and BTBR+ models (Error! Reference source not found.**G**). In the *Anks1b* Het model, Erg inhibition in both biological replicates, along with Nfe2l2 activation, predicted the downregulation of cell motility, neuronal apoptosis, and brain formation (Figure 5**G**, Error! Reference source not found.**H**). Overall effects were mild, similar to the Shank3* model, but reduced social behavior and increased seizure activity were predicted by activated Hif1a and inhibited Ptf1a (Anks1b Het2, Figure 5**H**). Although Erg inhibition was in opposition to the Fmr1 KO, inhibited Kdm5a and Nr4a1 in the Anks1b Het were similar to the Fmr1 KO (Figure 4**B**) and other models in Experiment 2 (Figure 2**D**), predicting altered energy metabolism at the synapse (Error! Reference source not found.**I**). In summary, networks of Regulatory Effects in each model show that common upstream regulators (Rest, Tlx3, Mecp2, Kdm5a) predict downstream effects on synaptic function and brain disorders.

**Figure 5.**
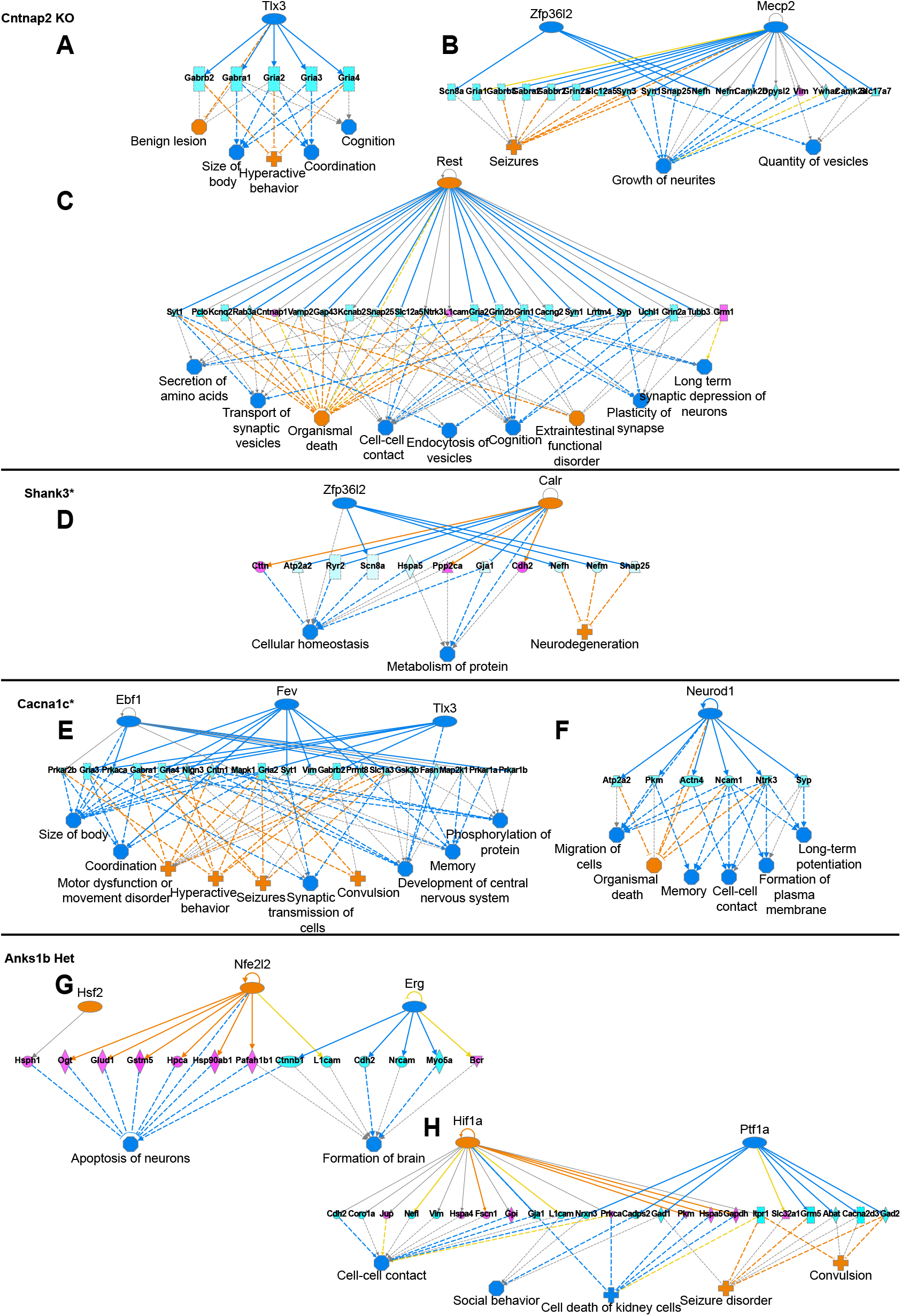
Altered synaptic composition predicts similar networks of functional effects in Cntnap2 KO, Shank3*, Cacna1c*, and Anks1b Het mice. **A-C**) Top regulatory networks for Cntnap2 KO mice. **D**) Top regulatory network for Shank3* mice. **E-F**) Top regulatory networks for Cacna1c* mice. **G**) Top regulatory networks for Anks1b Het mice. Upstream regulators and downstream effects were predicted in IPA from fold-changes of proteins in the postsynaptic proteome.

### 3.4 Synaptic changes predict altered Rho GTPase signaling across models

To identify cellular processes commonly altered in mouse models of autism, we compared activation of Canonical Pathways in IPA. In this domain, the relationships between ASD models were less consistent than in Upstream Analysis and Downstream Effects. In both Experiment 1 and Experiment 2, Analysis Match of Canonical Pathways revealed that some comparisons did not yield a similarity score (*p*>0.01), indicating that the relationship between them was neither significantly opposite nor similar (Kramer, et al. 2014). However, we did find similarity in the activation of molecular pathways between Fmr1 KO and Pten Het models, and among BTBR+, Cntnap2 KO, and Cacna1c* models in Experiment 1 (Figure 6**A**). In Experiment 2, we found that BTBR+, Cntnap2 KO, and Anks1b Het models were similar to each other (Figure 6**B**). In both experiments, significant changes (*z*-score > |2|) in pathways related to Rho GTPases featured prominently, including Signaling by Rho Family GTPases, RhoGDI Signaling, Rac Signaling, RhoA Signaling, Cdc42 Signaling, and Regulation of Actin-based Motility by Rho (Figure 6**A-B**). Directionality was largely consistent among mouse models, predicting activated signaling by Rho GTPases and inhibited regulation by RhoGDI, with the exception of Cacna1c* and Anks1b Het models (Figure 6**A-B**). Rho GTPase signaling was among the most enriched signaling pathways in the synaptic proteomes by STRING analysis (Figure 1**D**). The Rho (Ras homology) GTPases are an essential family of small guanine nucleoside triphosphate (GTP) binding proteins that hydrolyze GTP to regulate numerous aspects of neuronal development and function (Niftullayev and Lamarche-Vane 2019; Zamboni et al. 2018b), including synaptic transmission and plasticity (Murakoshi, Wang, and Yasuda 2011; Hedrick and Yasuda 2017).

**Figure 6.**
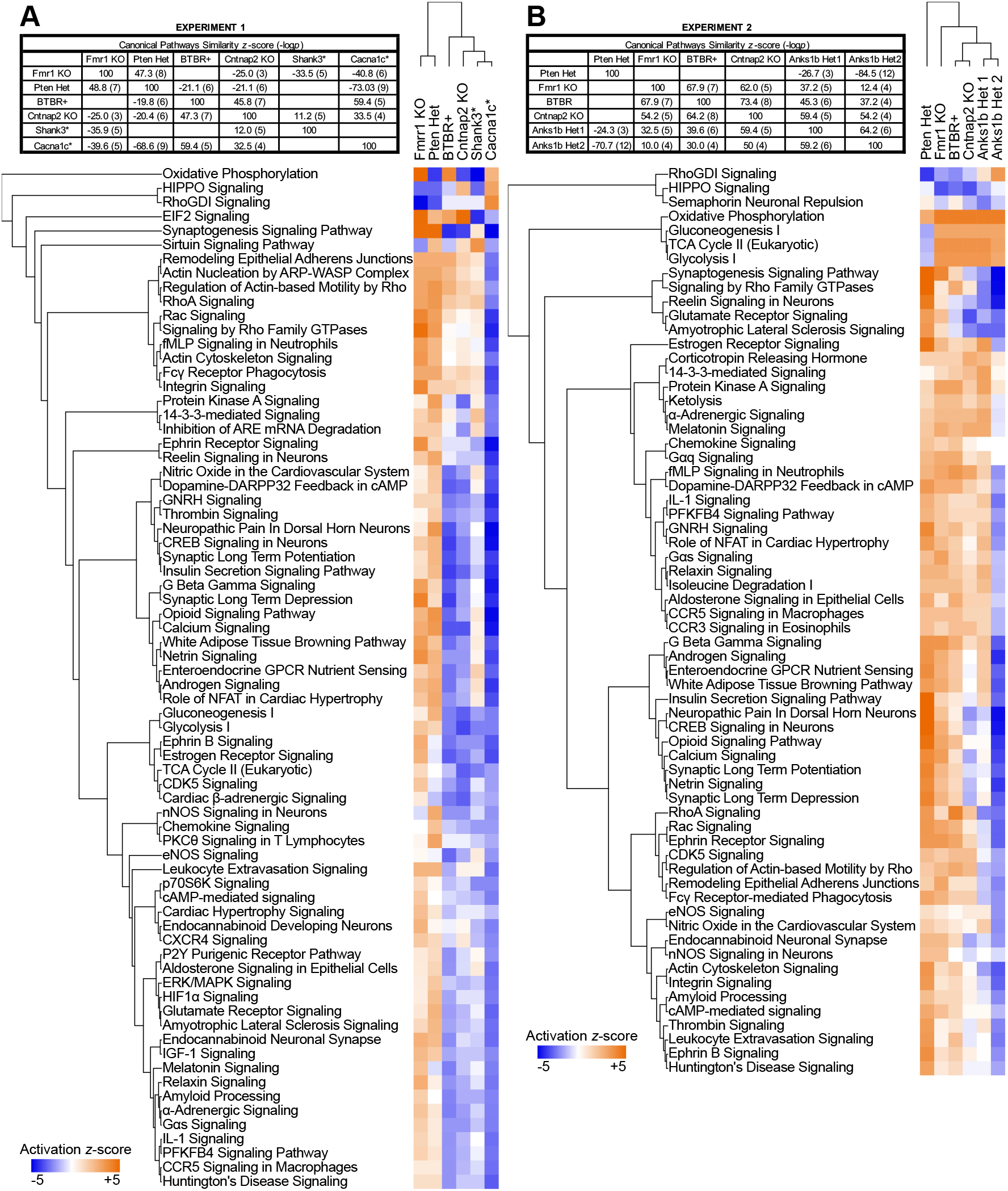
Predicted effects on Canonical Pathways show a distinct pattern of similarities and differences among mouse models of autism. **A**) For Canonical Pathways, similarity scores among ASD mouse models in Experiment 1 and Experiment 2 show similarities between the Fmr1 KO and Pten Het models, as opposed to the BTBR+, Cntnap2 KO, Shank3*, Cacna1c*, and Anks1b Het models (**B**). Clustering of ASD mouse models shows relationships among mouse models in Experiment 1 and Experiment 2. Tree diagrams show hierarchical cluster analysis of Canonical Pathways in IPA with activation *z*-scores >|2| and enrichment −log_10_(*p*-values) >5 (Benjamin-Hochberg corrected for multiple comparisons).

Since Rho GTPases have crucial roles in synaptic function that have been investigated in both human-based experiments and mouse models of ASD (Guo, Yang, and Shi 2020), we prioritized these pathways for further study. In hippocampal PSD fractions, only the small GTPase Rac1 met peptide cutoff criteria for inclusion in the synaptic proteome for both experiments. Analysis of the Rac Signaling pathway for each ASD mouse model shows strong activation in the Fmr1 KO (*z*-scores 3.96 and 2.98), Pten Het (*z*-scores 3.13 and 3.44), and BTBR+ strain (*z*-scores 1.04 and 2.52); weak activation in the Cntnap2 KO (*z*-scores 0.21 and 0.69) and Shank3* model (*z*-score 0.626); and inhibition of the Rac pathway in the Cacna1c* (*z*-score −3.96) and Anks1b Het models (*z*-score −0.69 and −2.98) (Table 1Error! Reference source not found.). In addition to Rac Signaling, RhoA and Cdc42 Signaling pathways displayed activation *z*-scores in a similar pattern: strong upregulation in Fmr1 KO, Pten Het, and BTBR+ strains, intermediate activation in Cntnap2 KO and Shank3*, and inhibition in Cacna1c* and Anks1b Het (Table 1). Depictions of the Rac Signaling pathway show that altered synaptic components of the pathway predicted activation in the Fmr1 KO (Experiment 1, Figure 7**A**) and BTBR+ models (Experiment 2, Figure 7**B**). In contrast, Rac Signaling was inhibited in the Cacna1c* (Figure 7**C**) and Anks1b models (Het2, Figure 7**D**). Overall, Rho-related signaling pathways showed opposing clusters of predicted activation (Fmr1 KO, Pten Het, BTBR+, Cntnap2 KO, Shank3*) and inhibition (Cacna1c* and Anks1b Het).

**Figure 7.**
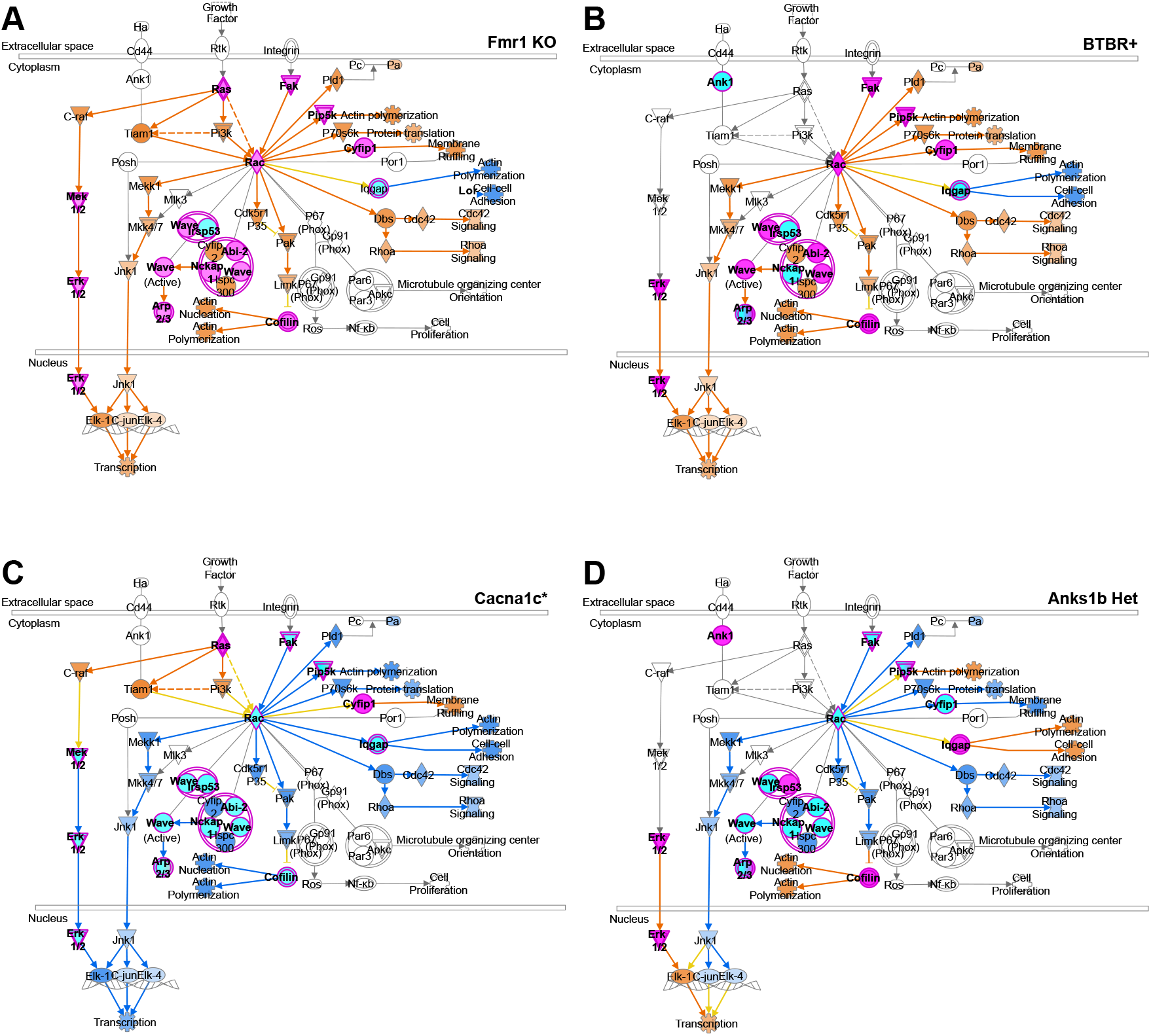
Rac Signaling Pathway is differentially regulated across mouse models of autism. Protein components of the Rac Signaling Pathway in IPA, are up- or downregulated in the postsynaptic proteomes of ASD mouse models, leading to predicted effects on pathway activation in **A**) Fmr1 KO mice (Experiment 1), **B**) BTBR+ mice (Experiment 2), **C**) Cacna1c* KO mice (Experiment 1), and **D**) Anks1b Het mice (Experiment 2). Purple outline = observed in proteome, Magenta = observed increase, Cyan = observed decrease, Orange arrow or fill = predicted activation, Blue arrow or fill = predicted inhibition, Yellow arrow = predicted effect inconsistent, Gray arrow or outline = effect not predicted, Solid line = direct effect, Dashed line = indirect effect.

### 3.5 Rho GTPases are differentially expressed in autism mouse models

For the mouse models of ASD showing significant upregulation of the Rac signaling pathway, with *z*-scores >2, Rac itself is upregulated (Rac in magenta, Figure 7**A,B**). To evaluate the synaptic expression of Rho family GTPases in mouse models for autism, we performed Western blots for Rac1, RhoA, and Cdc42 in synaptic fractions. Although we did not detect an immunoblot signal for RhoA and Cdc42 in PSD fractions using several antibodies, we were able to quantify Rac1 expression (Figure 8**A**). Consistent with the proteomics results showing strongest activation of the Rac signaling pathway, Rac1 expression was increased in the *Fmr1* KO and *Pten* Het models. However, Rac1 was also increased the *Anks1b* Het and no difference was measured in the BTBR+ strain or other models (Figure 8**B**). To determine whether the upregulation of Rac1 was specific to the synaptic compartment, we compared Rac1 expression levels in total lysate among all models. In contrast to our findings in the PSD fraction, we found broad reduction in Rac1 expression across ASD mouse models, including the Fmr1 KO, Pten Het, BTBR+ and Shank3* models in which Rac pathway activation was predicted in IPA (Figure 8**C**). The Anks1b Het model, in which inhibition of Rac signaling was predicted, also showed reduction of Rac1 in total lysate. Notably, Rac1 expression was significantly increased in the Cacna1c* model, which showed the strongest inhibition of Rac signaling as predicted from the proteomic data (Figure 6**A**).

**Figure 8.**
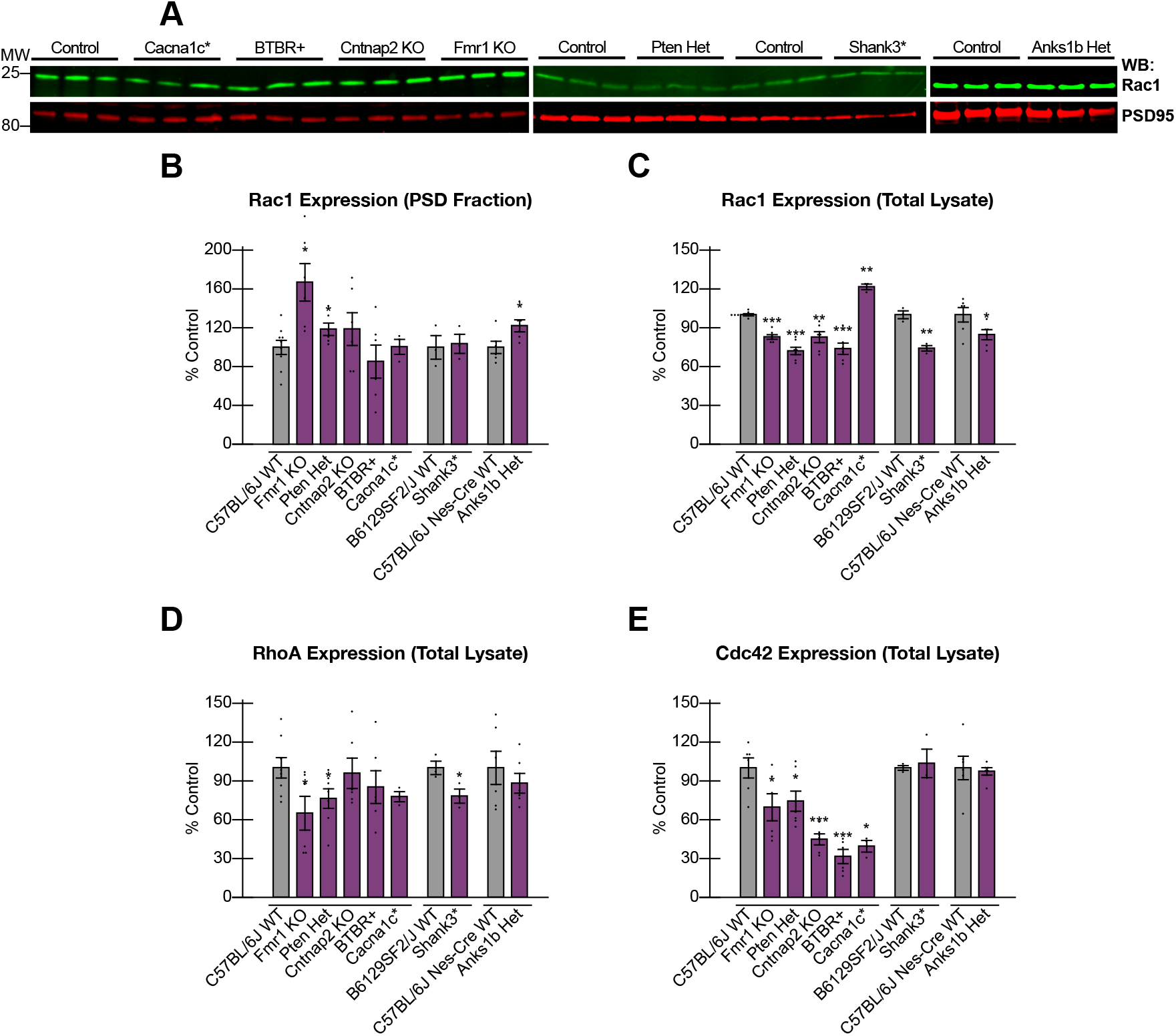
Rac1 is differentially expressed in synaptic fractions, and other Rho GTPases in total lysate, across mouse models of autism. **A**) Expression of Rac1, the only Rho GTPase detected in synaptic fractions, was (**B**) increased in Fmr1 KO, Pten Het, and Anks1b Het mice by Western blot (20 μg synaptic fraction). **C**) Expression of Rac1 in total lysate was reduced in all mouse models except the Cacna1c* model, where it was increased. **D**) Expression of RhoA in total lysate was reduced for Fmr1 KO, Pten Het, and Shank3* mice only. **E**) Expression of Cdc42 in total lysate was reduced for all models except Shank3* and Anks1b Het models, where no significant change was observed (30 μg total lysate). Bar graphs show mean ± SEM, *n*=3-10 samples for each mouse model and wild-type control strain, Student’s *t*-test, \**p*<0.05 \*\**p*<0.01 \*\*\**p*<0.001 as in **Error! Reference source not found.**.

In general, Rac pathway activation predicted in IPA was correlated with increased Rac1 expression in the PSD (Fmr1 KO and Pten Het) and reduced Rac1 in total lysate (Fmr1 KO, Pten Het, BTBR+, Cntnap2 KO, and Shank3*), while strong inhibition of Rac signaling was associated with increased Rac1 expression in total lysate (Cacna1c*). To test whether other Rho GTPases display a similar correlation between activation and expression, we performed Western blots for RhoA and Cdc42 in total lysate. RhoA expression was reduced in the Fmr1 KO, Pten Het, and Shank3* mouse models (Figure 8**C**). In agreement with our findings of reduced Rac1 expression, these models shared a predicted activation for the RhoA signaling pathway: Fmr1 KO *z*-scores 3.00 and 1.26, Pten Het *z*-scores 3.40 and 3.41, and Shank3* *z*-score 1.80 (Table 1). However, the trend with RhoA was less consistent than Rac1, with no significant change observed in other models with strong RhoA activation (BTBR+ *z*-scores 1.80 and 4.13, Cntnap2 KO *z*-scores 1.40 and 1.98) or inhibition (Cacna1c* *z*-score −2.60, Anks1b Het *z*-scores −2.34 and −2.69) (Figure 8**C**). For Cdc42, expression was markedly reduced across all mouse models except the Shank3* and Anks1b Het mice (Figure 8**D**). Accordingly, the Cdc42 signaling pathway was predicted to be activated in the Fmr1 KO (*z*-scores 2.36 and 3.00), Pten Het (*z*-scores 3.77 and 1.50), BTBR+ (*z*-scores 2.36 and 1.5), and Cntnap2 KO (*z*-scores 2.36 and 1.00).

However, the Cacna1c* model showed predicted inhibition of Cdc42 signaling (*z*-score −2.36), as of the other Rho GTPases. Although they showed no change in total Cdc42 expression, predicted changes for Cdc42 activation were weaker in Shank3* (*z*-score 0.94) and Anks1b Het models (*z*-scores −0.50 and −1.50) compared to other Rho GTPases in these mice (Table 1). Overall, the correlation between reduced total Cdc42 expression (by Western blot) with strong synaptic Cdc42 activation (in IPA) reflects the pattern in Rac1 and RhoA across mouse models. While several ASD models associated with synaptic Rho GTPase activation show global downregulation of GTPase expression (Fmr1 KO, Pten Het, BTBR+, Cntnap2 KO, Shank3*), others with predicted Rho inhibition demonstrate global upregulation (Cacna1c* and Anks1b Het).

## 4 Discussion

Here we compared the hippocampal postsynaptic proteomes from 7 mouse models for autism using quantitative mass spectrometry with isobaric tags (Figure 1), generating a resource for each model that can be mined for functional insights (Supplementary Data 1). Regulatory networks predicted by synaptic changes in each model reveal both novel results and patterns consistent with the literature (Figure 4, Figure 5, **Error! Reference source not found.**). In the Fmr1 KO model, regulatory effects showed that most proteins were upregulated (Figure 4**A-B**), which is consistent with a proteomic analysis of cortical synapses across the lifespan in *Fmr1* knockout mice (Tang et al. 2015). In the model of constitutive Pten haploinsufficiency, synaptic dysfunction (Figure 4**C-D**) is likely due to neuronal *Pten*, since altered synaptic composition was also observed in a neuron-specific conditional knockout mouse that displayed changes in activity patterns and repetitive behavior (Lugo et al. 2014). In the Cntnap2 KO mouse, the observed changes predicting increased seizures (Figure 5**B, Error! Reference source not found.E**) are consistent with clinical features of cortical dysplasia-focal epilepsy syndrome (Smogavec et al. 2016). Differences in the Shank3* model were milder than those observed in a previous study of the synaptic proteome in a *Shank3* knockout mouse (Reim, et al. 2017), although many different *Shank3* models have been developed that show distinct phenotypes (Varghese, et al. 2017). The *Shank3* InsG3680 mutation examined in this study models a human *SHANK3* mutation from a patient with autism, as opposed to other variants associated with schizophrenia and intellectual disability phenotypes in Phelan McDermid syndrome (Zhou, et al. 2016). For the Cacna1c* model, networks affecting heart failure and ion transport (**Error! Reference source not found.G**) were expected since Timothy syndrome is defined by calcium channel dysfunction leading to cardiac defects (Bader, et al. 2011). Regulatory effects in the Anks1b Het model showed subtle changes, with some similarities to Fmr1 KO and Pten Het models (Kdm5a inhibition, **Error! Reference source not found.I**) and some differences (Erg inhibition, Figure 5**G**). Predicted downregulation of synaptic genes by inhibited Erg and Ptf1a (Figure 5**G, Error! Reference source not found.H**) is relevant to *Anks1b* function, since its encoded protein AIDA-1 is a synapse-to-nucleus messenger that activates protein synthesis in response to synaptic activity (Jordan, et al. 2007). For the BTBR+ model, inhibited Fev and Neurod1 were shared with the Cacna1c* model, although BTBR+ mice shared the most upstream gene regulators with the Fmr1 KO and Pten Het models overall (Figure 2**C-D**).

Importantly, we directly compared synaptic proteomes in parallel, a necessary approach for finding shared phenotypes and broadly applicable therapeutic targets (Sestan and State 2018; Silverman and Ellegood 2018). Drawn from internally controlled measurements, our results demonstrate quantifiable differences in synaptic composition that support the presence of molecular subtypes in ASD rather than universal changes in the same direction. Using similarity *z-*scores and hierarchical clustering in IPA, we show that the Fmr1 KO, Pten Het models share predicted effects on upstream regulators (Figure 2**C-D**), diseases and functions (Figure 3), and cellular pathways (Figure 6). These results are consistent with the clinical features of the respective syndromes. In humans, macrocephaly has been observed in both Fragile X and PTEN hamartoma tumor syndromes (Butler et al. 2005; Chiu et al. 2007). Previous studies have shown that *Fmr1* and *Pten* mouse models share behavioral traits (Binder and Lugo 2017), transcriptional profiles (Lanz et al. 2013), and altered PI3K/AKT/mTOR signaling that affects proteostasis (Gross et al. 2019; Huber et al. 2015; Richter, Bassell, and Klann 2015). Here we report divergent activation signatures between the Fmr1 KO and Shank3* mutant, which was more similar to the Cacna1c* and Cntnap2 KO models overall (Figure 2). Recent comparison of synaptic interactomes and neuroanatomical changes among ASD mouse models showed that *Shank3* and *Fmr1* loss had similar effects, although these were found in other brain regions and *Shank3* deletion models (Brown, et al. 2018; Ellegood et al. 2015). Similarity between Shank3* and Cacna1c* mice is consistent with a direct behavioral comparison showing that *Shank3* and *Cacna1c* models share hypoactivity and anxiety phenotypes (Kabitzke, et al. 2018). Abnormal social and grooming behaviors are also shared between *Shank3* deletion and *Cntnap2* knockout models (Kabitzke et al. 2020), along with deficits in cerebellar motor learning (Kloth et al. 2015) and adult hippocampal neurogenesis (Cope et al. 2016).

Intermediate between the *Fmr1*/*Pten* mice and the *Shank3*/*Cntnap2*/*Cacna1c* models, the BTBR+ strain shows consistent similarities to Fmr1 KO and Pten Het in upstream regulators in both experiments (Figure 2**C-D**). As commonly studied models of syndromic and idiopathic ASD, Fmr1 KO and BTBR+ mice both respond to mGluR5 antagonism (Silverman et al. 2010), endocannabinoid potentiation (Wei et al. 2016), and intranasal dopamine (Chao et al. 2020), suggesting that changes in synaptic gene regulation can predict response to ASD treatment. Interestingly, BTBR+ and Shank3* models were only similar at the downstream level (Figure 3**A**), consistent with shared social phenotypes in BTBR+ and *Shank3* deletion mice (Rein, Yan, and Wang 2020). Divergent gene regulation and cellular pathways at hippocampal synapses suggests that behavioral similarities in BTBR+ and *Shank3* models are due to shared changes in the hippocampal cytosolic fraction (Daimon et al. 2015) or cortical interneurons (Gogolla et al. 2014). We show here that our *Anks1b* haploinsufficiency model is dissimilar to Fmr1 KO and Pten Het mice and similar to the Cntnap2 KO model across domains (Figure 2**B,D,** Figure 3**B,** Figure 6**B**). This pattern is similar to the Shank3* and Cacna1c* models, although they were not directly compared to Anks1b Het mice. Interestingly, these genes all regulate synaptic formation and ion channel function, supporting a role for AIDA-1 in activity-dependent membrane composition (Carbonell, et al. 2019; Tindi, et al. 2015) and structural changes at the synapse (Li et al. 2016). Overall, our results refine and extend previous reports of the phenotypic similarities among mouse models for autism and predict that common regulators contribute to synaptic dysfunction and ASD-related phenotypes.

In the 7 autism mouse models chosen for study, changes in the synaptic proteome converged on Rho family small GTPase signaling as a commonly altered cellular pathway (Figure 6). These results complement a growing body of evidence showing that altered activation of Rho GTPases is an important mechanism of disease in autism and other neurodevelopmental disorders (Pinto et al. 2010; Zeidan-Chulia et al. 2013; Guo, Yang, and Shi 2020). The Rho GTPases Rac1, RhoA, and Cdc42 act as molecular switches to coordinate cellular activities at neuronal membranes, including actin-based cytoskeletal remodeling (Niftullayev and Lamarche-Vane 2019). Their activation state is positively regulated by Rho guanine nucleotide exchange factors (RhoGEFs) and negatively by Rho GTPase activating proteins (RhoGAPs). Additionally, Rho GDP dissociation inhibitors (RhoGDIs) prevent Rho GTPase activation by sequestering them in the cytosol (Reichova et al. 2018). Consistent with our findings that synaptic proteins in the Rac pathway (Figure 7) and Rac1 itself is differentially regulated in ASD mouse models (Figure 8), evidence continues to emerge that Rac1 signaling plays an important role in the pathobiology of ASD and other neurodevelopmental disorders. Rac1 mutations associated with intellectual disability impair synaptic plasticity (Tian et al. 2018). Variants in the RhoGEF *TRIO* can cause ASD, intellectual disability, schizophrenia, or macrocephaly, with severe phenotypes predicted by Rac1 overactivation (Barbosa et al. 2020; Sadybekov et al. 2017; Katrancha et al. 2017). Mutations in the Rac1 RhoGEF *DOCK4* disrupts dendritic spine dynamics and synaptic transmission, leading to ASD, schizophrenia, and dyslexia (Huang et al. 2019; Xiao et al. 2013; Guo et al. 2019). Rac1 overactivation through loss of RhoGAPs can cause impaired dendritic development and cognition (*ARHGAP15*) (Zamboni et al. 2018a); X-linked intellectual disability (*OPHN1*) (Busti et al. 2020); and deficits in both axonal and dendritic development (*SRGAP*s) (Fossati et al. 2016; Perez, Sawmiller, and Tan 2016). Downstream of Rac1 in actin polymerization, mutations in *PAK*s and *WASF1* can cause macrocephaly, seizures, intellectual disability, or ASD (Horn et al. 2019; Ito et al. 2018).

Aside from genetic defects in its regulators and effectors, Rho GTPase pathways shows promise as a point of mechanistic convergence for other models of autism. Downstream of *Fmr1*, Rac1 overactivation modulates Cyfip1 regulation of dendritic spines and presynaptic morphology (De Rubeis et al. 2013; Hsiao et al. 2016). Neuroligins and neurexins, ASD-associated cell adhesion molecules related to *Cntnap2*, also converge on Cyfip1 function (Baudouin et al. 2012; Sledziowska, Galloway, and Baudouin 2019). Shank3 interacts with the RhoGAP Rich2 at the synapse to regulate AMPA receptor recycling and synaptic formation; loss of Rich2 leads to motor dysfunction, stereotypic behavior, and neophobia associated with Rac1 disinhibition (Raynaud et al. 2013; Sarowar et al. 2017; Sarowar et al. 2016). Shank3 also regulates NMDA receptor activity through cytoskeletal processes dependent on Rac1 (Duffney et al. 2013; Duffney et al. 2015). The ASD-related syndrome caused by 16p11.2 copy number variations seems to converge on RhoA, which can be upregulated by loss of *Kctd13* (Escamilla et al. 2017; Lin et al. 2015) or downregulated by loss of *Taok2* (Richter et al. 2019). In Timothy syndrome, *Cacna1c* mutations cause dendritic defects related to overactivation of RhoA (Krey et al. 2013). ASD-related variants in the *Stxbp5* gene lead to reduced dendritic branching and spine formation, along with reduced synaptic transmission attributed to upregulation of RhoA signaling (Shen et al. 2020; Cukier et al. 2014). Overall, Rac and RhoA pathways are commonly dysregulated in ASD and other neurodevelopmental disorders, and are therefore attractive targets for pharmacotherapy (Guo, Yang, and Shi 2020; Zamboni, et al. 2018b).

There are a number of caveats to our work. Multiple ASD models and control samples limited the number of simultaneous comparisons in 10-plex TMT-MS. Other models were excluded, including prominent genetic causes of ASD (*MECP2, UBE3A, TSC1/2, SYNGAP1, NLGN*/*NRXN*) (Hulbert and Jiang 2016; Verma, et al. 2019) and environmental risk factors (valproic acid, maternal immune activation) (Kazdoba, Leach, and Crawley 2016; Varghese, et al. 2017). However, these models affect similar pathways: *Fmr1* and *Ube3a* regulate proteostasis, *Pten* and *Tsc1*/*Tsc2* modulate PI3K/AKT/mTOR signaling, and *Cntnap2* is a member of the *Nrxn* family (Bagni and Zukin 2019). *Anks1b* encodes the synaptic adaptor protein AIDA-1, which binds Shank3 and Syngap1 and behaves similarly as an effector of long-term potentiation (Carbonell, et al. 2019; Dosemeci et al. 2016; Li, et al. 2016; Tindi, et al. 2015). Proteomic studies of maternal immune activation also converge on synaptic dysfunction (Lombardo et al. 2018; Gyorffy et al. 2016). Comparing models at a single developmental timepoint and brain region limits the scope of observed changes, since models of ASD show behavioral and biological phenotypes that differ based on the spatiotemporal targeting of the gene (Del Pino, Rico, and Marin 2018). A proteomic study of cortical synapses noted smaller differences in Fmr1 KO mice after 3 weeks of age (Tang, et al. 2015), and both Shank3 and Syngap1 show changes in interaction partners throughout development (Li et al. 2017; Li, et al. 2016). However, young adulthood in mice is a developmental stage that can be effectively targeted through genetic and pharmacological interventions (Yan et al. 2018; Duffney, et al. 2015). Since ASD is diagnosed in early childhood, after critical periods of fetal brain development, identifying and potentially correcting molecular phenotypes that have already developed has translational value. Although hippocampal circuits have been extensively studied for their robust expression of synaptic plasticity and its implications for cognitive function, projection neurons in the neocortex, striatum, and cerebellum are also relevant to ASD animal models (Golden, Buxbaum, and De Rubeis 2018). Fractionation of the PSD limits our findings to synaptic profiles in glutamatergic neurons, but ASD risk genes also influence interneuron development (*Cntnap2*) (Penagarikano, et al. 2011) and oligodendrocyte maturation (*Pten*) (Lee et al. 2019). Importantly, the Rho GTPases that regulate synaptic changes in ASD models (Figure 6) are expressed in brain regions and cell types relevant to ASD: increased expression of Rac1 was found in cortical tissue from autism patients (Fatemi et al. 2013), Rac overactivation affects development of inhibitory hippocampal interneurons (Zamboni et al. 2016), and Rac1 is involved in development of oligodendrocytes and cerebellum, potentially contributing to impaired connectivity (Zeidan-Chulia et al. 2016). Future experiments to compare proteomes across cell types and developmental stages could confirm common synaptic mechanisms in brain regions, quantify relationships among monogenic ASDs, and refine molecular subtypes of autism (Sestan and State 2018; Stessman, Turner, and Eichler 2016; Iakoucheva, Muotri, and Sebat 2019).

Since many important risk factors for ASD converge at the synapse, investigating shared changes at the proteomic level can yield mechanistic insights and therapeutic targets. We propose that synaptic proteomes, as quantifiable phenotypes of diverse ASD models, capture the combined influences of genetic, epigenetic, transcriptomic, translational, and degradative factors implicated in ASD etiology. By focusing on the proteome, we bypass the need to understand the complex or unclear functional outcomes of these multiple factors, allowing us to directly measure their net results at the synapse. Using TMTs and quantitative proteomic methods, we found functional similarities in the postsynaptic profiles of mouse models for monogenic and idiopathic ASD. We identified Rho GTPases, especially Rac1 signaling, as crucial cellular pathways disrupted in the hippocampus across ASD mouse models. These results demonstrate that parallel processing of samples from models targeting multiple ASD risk genes can yield convergent results. They also provide a resource to supplement ongoing work in widely studied ASD mouse models and generate hypotheses for further investigation. We expect that validating Rho GTPase signaling as a point of convergence in mouse models of autism will help clarify the pathogenesis of ASD, and that future studies will pursue key nodes in the Rac1 signaling cascade as potential therapeutic targets.

## Acknowledgements

Supported by NIH R01AG039521 and NIH R56MH115201 to BAJ, T32GM007288 to AUC, NIH S10RR027990 to TAN, and NIH R01MH108519 to DTP. Significant support for this work came from the Rose F. Kennedy Intellectual and Developmental Disabilities Research Center (IDDRC), which is funded through the center grant NIH U54HD090260.

