## Supplemental tables and figures for "Comparing synaptic proteomes across seven mouse models for autism reveals molecular subtypes and deficits in Rho GTPase signaling"

**Supplementary Table 1. Hippocampal postsynaptic proteome from mouse models of autism (Experiment 1).**

Proteins quantified by >2 peptides in each sample of postsynaptic density (PSD) enriched fractions from ASD mouse models in Experiment 1 (788 proteins).

| | | | | | | | | | |
|---|---|---|---|---|---|---|---|---|---|
| Aak1 | Atp2b1 | Clasp1 | Dpysl5 | Grm1 | Lsamp | Nefl | Prkar1a | Rtn4 | Syngr3 |
| Abcd3 | Atp2b2 | Clasp2 | Dsp | Grm2 | Lzts3 | Nefm | Prkar1b | Ryr2 | Synj1 |
| Abhd12 | Atp5a1 | Cldn11 | Dst | Grm3 | Macf1 | Negr1 | Prkar2a | Samm50 | Synpo |
| Abi1 | Atp5b | Clmn | Dync1h1 | Grm5 | Madd | Nf1 | Prkar2b | Sbf1 | Syp |
| Abi2 | Atp5c1 | Clta | Dync1li1 | Grm7 | Magi2 | Nfasc | Prkcb | Scai | Syt1 |
| Ablim1 | Atp5h | Cltc | Eef1a1 | Gsk3b | Map1a | Nipsnap1 | Prkce | Scn8a | Syt7 |
| Ablim2 | Atp5l | Cnksr2 | Eef1a2 | Gucy1b1 | Map1b | Nipsnap2 | Prkcg | Scn9a | Tanc1 |
| Abr | Atp5o | Cnp | Eef1g | H2afx | Map2 | Nlgn1 | Prmt8 | Sept11 | Tanc2 |
| Acat1 | Atp5pb | Cntn1 | Eef2 | H3f3a | Map2k1 | Nlgn2 | Prnp | Sept2 | Tbc1d10b |
| Acly | Atp6ap1 | Cntnap1 | Efr3a | Hacd3 | Map4 | Nlgn3 | Prr7 | Sept3 | Tbk1 |
| Aco2 | Atp6v0a1 | Coro1a | Efr3b | Hadha | Map4k4 | Nme1 | Prrt1 | Sept4 | Tecr |
| Acot9 | Atp6v0d1 | Coro1c | Elfn1 | Hadhb | Map6 | Nos1 | Prxl2a | Sept5 | Tenm2 |
| Acta1 | Atp6v1a | Coro2b | Elfn2 | Hap1 | Map6d1 | Nptn | Psd | Sept6 | Tenm3 |
| Actg1 | Atp6v1b2 | Cox4i1 | Elmo2 | Hapln1 | Mapk1 | Nrcam | Psd3 | Sept7 | Tenm4 |
| Actn1 | Atp6v1c1 | Cpeb3 | Emc2 | Hba1 | Mapre2 | Nrxn1 | Ptk2b | Sept8 | Tfam |
| Actn2 | Atp6v1d | Cpne6 | Enah | Hexb | Mapre3 | Nrxn3 | Ptprd | Sept9 | Thy1 |
| Actn4 | Atp6v1e1 | Crmp1 | Eno1 | Hist1h2bc | Mapt | Nsf | Ptprs | Sfn | Tjp1 |
| Actr1a | Atp6v1g2 | Cs | Eno2 | Hist1h4a | Marc2 | Ntm | Pura | Sfxn3 | Tln2 |
| Actr1b | Atp6v1h | Csnk1a1 | Epb41l1 | Hk1 | Mark1 | Ntrk3 | Pygb | Sfxn5 | Tmem121b |
| Actr2 | Auh | Csnk1d | Epb41l2 | Hnrnpa2b1 | Mark2 | Nwd2 | Rab10 | Sgip1 | Tmem33 |
| Actr3 | Baiap2 | Csnk1e | Epb41l3 | Homer1 | Mark3 | Ogdh | Rab11fip5 | Sh3gl2 | Tmod2 |
| Actr3b | Basp1 | Csnk1g2 | Erbin | Homer2 | Mark4 | Ogt | Rab1a | Sh3kbp1 | Tmx2 |
| Adam11 | Bcas1 | Csnk2a1 | Erc1 | Homer3 | Mast3 | Olfm1 | Rab35 | Shank1 | Tnik |
| Adam22 | Bcr | Csnk2a2 | Erc2 | Hsd17b4 | Mbp | Olfm2 | Rab3a | Shank2 | Tpi1 |
| Adcy1 | Bdh1 | Csnk2b | Evl | Hsp90aa1 | Mdh1 | Omg | Rac1 | Shank3 | Tppp |
| Add1 | Begain | Csrp1 | Ezr | Hsp90ab1 | Mdh2 | Opa1 | Rap1b | Shisa6 | Traf3 |
| Add2 | Bin1 | Ctnna2 | Fam81a | Hspa12a | Mgst3 | Oxct1 | Rapgef2 | Shisa7 | Trim3 |
| Add3 | Braf | Ctnnb1 | Farp1 | Hspa1a | Mink1 | Pacs1 | Rapgef4 | Sik3 | Trio |
| Adgrb1 | Brinp1 | Ctnnd1 | Fasn | Hspa2 | Mlf2 | Pafah1b1 | Rasal1 | Sipa1l1 | Ttc7b |
| Adgrb2 | Brsk1 | Ctnnd2 | Fbxo2 | Hspa5 | Mog | Palm | Rasgrf1 | Sipa1l3 | Tuba1a |
| Adgrb3 | Brsk2 | Cttn | Fbxo41 | Hspa8 | Mpc2 | Palm2 | Rgs7 | Slc12a5 | Tuba1c |
| Adgrl1 | Bsn | Cttnbp2 | Fech | Hspa9 | Mpp2 | Pclo | Rimbp2 | Slc17a7 | Tuba4a |
| Adgrl3 | Ca4 | Cul3 | Flot1 | Hspd1 | Mprip | Pde2a | Rims1 | Slc1a2 | Tubb2a |
| Afdn | Cacna1a | Cycs | Flot2 | Htt | Mrtfb | Pde4d | Rims2 | Slc1a3 | Tubb3 |
| Agap2 | Cacna1b | Cyfip1 | Fry | Icam5 | MT-CO2 | Pdha1 | Rogdi | Slc25a1 | Tubb4a |
| Agap3 | Cacna1e | Cyfip2 | Gabarapl2 | Idh2 | MT-ND1 | Pdhb | Rp2 | Slc25a11 | Tubb4b |
| Ahsa1 | Cacna2d1 | Cyld | Gabbr1 | Idh3g | Mtch1 | Pdhx | Rph3a | Slc25a12 | Tubb5 |
| Ajm1 | Cacna2d3 | Cyp46a1 | Gabbr2 | Igsf21 | Mtch2 | Peak1 | Rpl10a | Slc25a22 | Tubb6 |
| Ak5 | Cacnb1 | Cyth2 | Gabra1 | Igsf8 | Mtmr1 | Pex5l | Rpl12 | Slc25a3 | Tufm |
| Akap5 | Cacnb2 | Dab2ip | Gabra2 | Ildr2 | Myh10 | Pfkl | Rpl13 | Slc25a4 | Uba52 |
| Aldoa | Cacnb3 | Dagla | Gabra5 | Immt | Myh14 | Pfkm | Rpl13a | Slc25a5 | Uchl1 |
| Aldoc | Cacnb4 | Dbn1 | Gabrb2 | Ina | Myh9 | Pfkp | Rpl14 | Slc2a1 | Unc13a |
| Alg2 | Cacng2 | Dclk1 | Gabrb3 | Iqcb1 | Myl12b | Pgam1 | Rpl15 | Slc4a10 | Uqcrc1 |
| Amph | Cacng3 | Dclk2 | Gabrg2 | Iqgap2 | Myl6 | Pgam5 | Rpl17 | Slc4a4 | Uqcrc2 |
| Ank2 | Cacng8 | Dctn1 | Gap43 | Iqsec1 | Myo18a | Pgk1 | Rpl21 | Slc4a7 | Usp5 |
| Ank3 | Cadps | Dctn2 | Gapdh | Iqsec2 | Myo1d | Phactr1 | Rpl23 | Slc4a8 | Usp9x |
| Anks1b | Calcoco1 | Ddn | Gatd3a | Iqsec3 | Myo5a | Phb | Rpl23a | Slc8a1 | Vamp2 |
| Ap1b1 | Calml3 | Ddx1 | Gda | Itpka | Myo6 | Phb2 | Rpl24 | Slc8a2 | Vapa |
| Ap2a1 | Camk2a | Ddx3y | Gdap1 | Jcad | Nap1l1 | Pi4ka | Rpl26 | Slitrk5 | Vapb |
| Ap2a2 | Camk2b | Dgkb | Gdi1 | Jph3 | Napb | Pip4k2b | Rpl27 | Snap25 | Vcpip1 |
| Ap2b1 | Camk2d | Dgkg | Gfap | Jup | Nav1 | Pip5k1c | Rpl28 | Snap47 | Vdac1 |
| Ap2m1 | Camk2g | Diaph2 | Git1 | Kalrn | Nbea | Pitpnm2 | Rpl3 | Snap91 | Vdac2 |
| Ap2s1 | Camkv | Dlat | Gja1 | Kcna2 | Ncald | Pkm | Rpl31 | Snd1 | Vdac3 |
| Ap3b2 | Camsap3 | Dld | Gjc2 | Kcnab1 | Ncam1 | Pkp4 | Rpl35a | Snph | Vim |
| Ap3d1 | Capn5 | Dlg1 | Glud1 | Kcnab2 | Ncan | Plec | Rpl4 | Sorbs1 | Vps45 |
| Arc | Capza2 | Dlg2 | Glul | Kcnd2 | Ncdn | Plekha6 | Rpl5 | Sorbs2 | Vsnl1 |
| Arf1 | Capzb | Dlg3 | Gnai1 | Kcnd3 | Nckap1 | Plp1 | Rpl6 | Specc1 | Wasf1 |
| Arhgap1 | Carmil1 | Dlg4 | Gnai2 | Kcnj4 | Nckipsd | Plpp3 | Rpl7 | Sptan1 | Wdr1 |
| Arhgap21 | Cask | Dlgap1 | Gnao1 | Kcnma1 | Ndufa10 | Plppr4 | Rpl7a | Sptb | Wdr37 |
| Arhgap32 | Caskin1 | Dlgap2 | Gnaq | Kcnq2 | Ndufa13 | Plxna1 | Rpl8 | Sptbn1 | Wdr47 |
| Arhgap39 | Ccdc177 | Dlgap3 | Gnas | Kctd12 | Ndufa2 | Ppfia2 | Rpl9 | Src | Wdr7 |
| Arhgap44 | Ccny | Dlgap4 | Gnaz | Kctd16 | Ndufa4 | Ppfia3 | Rplp0 | Srcin1 | Wfs1 |
| Arhgef17 | Cd47 | Dlst | Gnb1 | Kiaa1217 | Ndufa6 | Ppia | Rpn1 | Srgap3 | Ywhab |
| Arhgef2 | Cdc42 | Dmxl2 | Got2 | Kiaa1549 | Ndufa7 | Ppib | Rpn2 | Strip1 | Ywhae |
| Arhgef7 | Cdh10 | Dnaja1 | Gphn | Kif2a | Ndufa9 | Ppp1ca | Rps10 | Strn | Ywhag |
| Arpc1a | Cdh11 | Dnaja2 | Gpi | Kif5c | Ndufb10 | Ppp1cc | Rps13 | Strn3 | Ywhah |
| Arpc2 | Cdh2 | Dnaja3 | Gpm6a | Klc2 | Ndufb4 | Ppp1r12a | Rps16 | Strn4 | Ywhaq |
| Arpc3 | Cdk5 | Dnajb1 | Gpr158 | Krt76 | Ndufb5 | Ppp1r9b | Rps18 | Stt3b | Ywhaz |
| Arpc4 | Cdkl5 | Dnajb6 | Gprin1 | Ksr1 | Ndufb8 | Ppp2ca | Rps19 | Stx1b | |
| Arpc5 | Cend1 | Dnajc11 | Grasp | Ktn1 | Ndufb9 | Ppp2r1a | Rps2 | Stxbp1 | |
| Arpc5l | Cep170 | Dnm1 | Gria1 | L1cam | Ndufs1 | Ppp2r5c | Rps23 | Stxbp5l | |
| Arvcf | Cep170b | Dnm1l | Gria2 | Lgi1 | Ndufs2 | Ppp2r5e | Rps3 | Sucla2 | |
| Astn1 | Cfl1 | Dnm2 | Gria3 | Lmtk3 | Ndufs3 | Ppp3ca | Rps3a | Suclg1 | |
| Atad3a | Chchd3 | Dnm3 | Gria4 | Lrrc4b | Ndufs4 | Ppp3cb | Rps4x | Sv2b | |
| Atat1 | Chchd6 | Dock3 | Grik2 | Lrrc59 | Ndufs8 | Prdx1 | Rps6 | Syn1 | |
| Atp1a1 | Cit | Dock4 | Grik5 | Lrrc7 | Ndufv1 | Prdx2 | Rps7 | Syn2 | |
| Atp1a2 | Ckap4 | Dock9 | Grin1 | Lrrc8a | Ndufv2 | Prdx5 | Rps9 | Syn3 | |
| Atp1a3 | Ckap5 | Dpp6 | Grin2a | Lrrtm1 | Nedd4 | Prickle2 | Rtcb | Syncrip | |
| Atp1b1 | Ckb | Dpysl2 | Grin2b | Lrrtm2 | Nedd4l | Prkaca | Rtn1 | Syne1 | |
| Atp2a2 | Ckmt1 | Dpysl3 | Grk3 | Lrrtm4 | Nefh | Prkacb | Rtn3 | Syngap1 | |

**Supplementary Table 2. Hippocampal postsynaptic proteome from mouse models of autism (Experiment 2).**

Proteins quantified by >9 peptides in each sample of postsynaptic density (PSD) enriched fractions from ASD mouse models in Experiment 2 (830 proteins).

| | | | | | | | | | | |
|---|---|---|---|---|---|---|---|---|---|---|
| Aak1 | Arhgef2 | Capzb | Dlgap1 | Gap43 | Inpp4a | Mdh1 | Pclo | Rapgef2 | Specc1 | Usp5 |
| Aars | Arhgef7 | Cask | Dlgap2 | Gapdh | Iqgap2 | Mdh2 | Pdcd6ip | Rapgef4 | Speg | Usp9x |
| Abat | Arpc1a | Caskin1 | Dlgap3 | Gas7 | Iqsec1 | Mical3 | Pde2a | Rasal1 | Sphkap | Vapa |
| Abcd3 | Arpc2 | Ccdc177 | Dlgap4 | Gda | Iqsec2 | Mink1 | Pde4a | Rasgrf1 | Sptan1 | Vcan |
| Abi1 | Arpc5 | Ccdc88a | Dlst | Gdap1 | Iqsec3 | Mobp | Pde4d | Rasgrf2 | Sptb | Vcl |
| Abi2 | Arpc5l | Cct2 | Dmtn | Gdi1 | Itpka | Mog | Pdha1 | Rdx | Sptbn1 | Vcp |
| Abl2 | Arvcf | Cct3 | Dmxl2 | Gdi2 | Itpr1 | Mpp2 | Pdhb | Rimbp2 | Srcin1 | Vcpip1 |
| Ablim1 | Asap1 | Cct4 | Dnaja1 | Gfap | Itsn1 | Mpp6 | Pdhx | Rims1 | Srgap2 | Vdac1 |
| Ablim2 | Astn1 | Cct6a | Dnaja2 | Git1 | Itsn2 | Mprip | Peak1 | Rims2 | Srgap3 | Vdac2 |
| Abr | Atad3a | Cct8 | Dnaja3 | Gja1 | Jup | Msn | Pex5l | Rmdn3 | Stim2 | Vdac3 |
| Acaa2 | Atat1 | Cdc42bpa | Dnajb6 | Gk | Kalrn | Mtch1 | Pfkl | Robo2 | Strn | Vim |
| Acat1 | Atp1a1 | Cdc42bpb | Dnajc11 | Glg1 | Kbtbd11 | Mtch2 | Pfkm | Rock2 | Strn3 | Vps35 |
| Acly | Atp1a2 | Cdh10 | Dnajc6 | Gls | Kcna1 | Mtdh | Pfkp | Rph3a | Strn4 | Vsnl1 |
| Aco2 | Atp1a3 | Cdh2 | Dnm1 | Glud1 | Kcna2 | Mthfd1 | Pgam5 | Rpl3 | Stx1a | Wasf1 |
| Acot7 | Atp1b1 | Cdkl5 | Dnm1l | Glul | Kcnab2 | Mtmr1 | Pgk1 | Rpl4 | Stx1b | Wdfy3 |
| Acsl6 | Atp2a2 | Cep170 | Dnm2 | Gnai2 | Kcnd2 | Mtor | Phactr1 | Rplp0 | Stxbp1 | Wdr1 |
| Actg1 | Atp2b1 | Cep170b | Dnm3 | Gnao1 | Kcnma1 | Mtus2 | Phb | Rpn1 | Sucla2 | Wdr37 |
| Actn1 | Atp2b2 | Cfl1 | Dock3 | Gnaq | Kcnq2 | Mycbp2 | Phb2 | Rps3 | Sv2a | Wdr47 |
| Actn2 | Atp2b4 | Chchd3 | Dock4 | Gnas | Kctd12 | Myh10 | Phf24 | Rps3a | Syn1 | Wdr7 |
| Actn4 | Atp5a1 | Chchd6 | Dock7 | Gnaz | Kctd16 | Myh14 | Pi4ka | Rrbp1 | Syn2 | Wfs1 |
| Actr1a | Atp5b | Cit | Dock9 | Gnb1 | Kiaa1107 | Myh9 | Pip4k2b | Rtn1 | Syn3 | Wipf2 |
| Actr1b | Atp5c1 | Ckap4 | Dpp10 | Gnb2 | Kiaa1109 | Myo18a | Pip5k1c | Rtn3 | Syne1 | Wipf3 |
| Actr2 | Atp5h | Ckap5 | Dpp6 | Gnb4 | Kiaa1211 | Myo1d | Pitpna | Rtn4 | Syngap1 | Wnk1 |
| Actr3 | Atp5pb | Ckb | Dpysl2 | Got1 | Kiaa1217 | Myo5a | Pitpnm1 | Ryr2 | Synj1 | Wnk2 |
| Actr3b | Atp6v0a1 | Ckmt1 | Dpysl3 | Got2 | Kiaa1468 | Myo6 | Pitpnm2 | Ryr3 | Synpo | Ywhab |
| Adam22 | Atp6v0d1 | Clasp1 | Dpysl4 | Gpd2 | Kiaa1549 | Napb | Pkm | Samm50 | Syt1 | Ywhae |
| Adam23 | Atp6v1a | Clasp2 | Dpysl5 | Gphn | Kif1a | Napg | Pkp4 | Sbf1 | Syt2 | Ywhag |
| Adcy9 | Atp6v1b2 | Clip1 | Dst | Gpi | Kif21a | Nav1 | Plcb1 | Scai | Syt7 | Ywhah |
| Add1 | Atp6v1c1 | Clip2 | Dtna | Gpr158 | Kif2a | Nav3 | Plec | Scn1a | Tagln3 | Ywhaq |
| Add2 | Atp6v1d | Clmn | Dync1h1 | Gprin1 | Kif3a | Nbea | Plekha6 | Sdha | Tanc1 | Ywhaz |
| Add3 | Atp6v1e1 | Cltc | Dync1i1 | Gria1 | Kif5a | Ncam1 | Plp1 | Sdhb | Tanc2 | |
| Adgrb1 | Atp6v1h | Cnksr2 | Dync1i2 | Gria2 | Kif5b | Ncam2 | Plppr4 | Septin11 | Taok1 | |
| Adgrb2 | Atp8a1 | Cnp | Dync1li1 | Gria3 | Kif5c | Ncan | Plxna1 | Septin3 | Tbc1d10b | |
| Adgrb3 | Atxn2l | Cntn1 | Eef1a1 | Gria4 | Klc1 | Ncdn | Plxna4 | Septin4 | Tbk1 | |
| Adgrl1 | Auh | Cntnap1 | Eef1a2 | Grin1 | Klc2 | Nckap1 | Ppfia2 | Septin5 | Tcp1 | |
| Adgrl3 | Baiap2 | Cntnap2 | Eef1g | Grin2a | Kpnb1 | Nckipsd | Ppfia3 | Septin6 | Tenm1 | |
| Afdn | Basp1 | Coro1a | Eef2 | Grin2b | Ksr1 | Ndufa10 | Ppia | Septin7 | Tenm2 | |
| Afg3l2 | Bcan | Coro1b | Efr3b | Grip1 | Ktn1 | Ndufa12 | Ppp1ca | Septin8 | Tenm3 | |
| Agap2 | Bcas1 | Coro1c | Eif4b | Grk2 | L1cam | Ndufa9 | Ppp1cb | Septin9 | Tenm4 | |
| Agap3 | Bcr | Coro2b | Eif4g1 | Grm1 | Larp1 | Ndufs1 | Ppp1cc | Sfxn3 | Tfam | |
| Ahcyl1 | Bdh1 | Cpne6 | Elfn1 | Grm2 | Lasp1 | Ndufs2 | Ppp1r12a | Sgip1 | Tjp1 | |
| Ahcyl2 | Begain | Crmp1 | Elfn2 | Grm3 | Ldha | Ndufv1 | Ppp1r13b | Sh3gl2 | Tjp2 | |
| Aifm1 | Bin1 | Crocc | Elmo2 | Grm5 | Ldhb | Ndufv2 | Ppp1r9b | Sh3kbp1 | Tkt | |
| Ajm1 | Braf | Cs | Enah | Grm7 | Letm1 | Nebl | Ppp2ca | Shank1 | Tln1 | |
| Ak5 | Brinp1 | Csnk1d | Eno1 | Gsk3a | Lgi1 | Nedd4l | Ppp2cb | Shank2 | Tln2 | |
| Akap2 | Brsk1 | Csnk2a1 | Eno2 | Gsk3b | Limch1 | Nefh | Ppp2r1a | Shank3 | Tmod2 | |
| Akap5 | Brsk2 | Csnk2a2 | Epb41l1 | Gstm1 | Lmna | Nefl | Ppp2r2a | Shisa6 | Tnik | |
| Aldh1l1 | Bsn | Ctnna1 | Epb41l2 | Hadha | Lmnb1 | Nefm | Ppp3ca | Shisa7 | Tnks1bp1 | |
| Aldh2 | C2cd2l | Ctnna2 | Epb41l3 | Hadhb | Lmnb2 | Negr1 | Ppp3cb | Sipa1l1 | Tnr | |
| Aldh5a1 | Cacna1a | Ctnnb1 | Epha4 | Hapln1 | Lmtk3 | Nf1 | Prdx5 | Sipa1l3 | Tomm70a | |
| Aldoa | Cacna1b | Ctnnd1 | Eps15l1 | Hcn1 | Lrpprc | Nfasc | Prickle2 | Skp1 | Tpi1 | |
| Aldoc | Cacna1e | Ctnnd2 | Erbin | Hecw2 | Lrrc59 | Nhsl2 | Prkaca | Slc12a2 | Tppp | |
| Amer2 | Cacna2d1 | Cttn | Erc1 | Hk1 | Lrrc7 | Nlgn2 | Prkar1a | Slc12a5 | Traf3 | |
| Ampd2 | Cacna2d3 | Cttnbp2 | Erc2 | Homer1 | Lrrc8a | Nlgn3 | Prkar1b | Slc17a7 | Trappc9 | |
| Amph | Cacnb1 | Cul3 | Erlin2 | Homer2 | Lsamp | Nlgn4l | Prkar2a | Slc1a2 | Trim3 | |
| Ank1 | Cacnb2 | Cyc1 | Evl | Homer3 | Lzts3 | Nos1 | Prkar2b | Slc1a3 | Trim46 | |
| Ank2 | Cacnb3 | Cyfip1 | Exoc2 | Hpca | Macf1 | Npepps | Prkca | Slc25a11 | Trim9 | |
| Ank3 | Cacnb4 | Cyfip2 | Exoc4 | Hsd17b4 | Madd | Nptn | Prkcb | Slc25a12 | Trio | |
| Anks1b | Cacng8 | Cyld | Exoc7 | Hsp90aa1 | Magi2 | Nrcam | Prkce | Slc25a18 | Tsc2 | |
| Anxa6 | Cadm2 | Daam1 | Exoc8 | Hsp90ab1 | Map1a | Nrxn1 | Prkcg | Slc25a22 | Ttbk1 | |
| Ap1b1 | Cadm3 | Dab2ip | Ezr | Hspa12a | Map1b | Nrxn3 | Prnp | Slc25a3 | Ttc7b | |
| Ap2a1 | Cadps | Dagla | Fam171a2 | Hspa1a | Map1s | Nsf | Prrc2a | Slc25a4 | Tuba1a | |
| Ap2a2 | Cadps2 | Dars | Fam81a | Hspa1b | Map2 | Nt5dc3 | Prrc2c | Slc25a46 | Tuba4a | |
| Ap2b1 | Calcoco1 | Dbn1 | Farp1 | Hspa2 | Map4 | Ntm | Prrt3 | Slc25a5 | Tubb2a | |
| Ap2m1 | Calml3 | Dbnl | Fasn | Hspa4 | Map4k4 | Numbl | Psd | Slc27a4 | Tubb3 | |
| Ap3b2 | Camk2a | Dclk1 | Fbxo41 | Hspa4l | Map6 | Nwd2 | Psd3 | Slc32a1 | Tubb4a | |
| Ap3d1 | Camk2b | Dclk2 | Fh | Hspa5 | Map6d1 | Ogdh | Psma1 | Slc3a2 | Tubb4b | |
| Apc | Camk2d | Dctn1 | Flot1 | Hspa8 | Map7d1 | Ogt | Ptk2b | Slc4a10 | Tubb5 | |
| Arfgap1 | Camk2g | Dctn2 | Flot2 | Hspa9 | Map7d2 | Opa1 | Ptprd | Slc4a4 | Tubb6 | |
| Arfgef2 | Camkv | Ddn | Fmn2 | Hspd1 | Mapk1 | Oxct1 | Ptprs | Slc6a11 | Tufm | |
| Arfgef3 | Camsap1 | Ddx3x | Fmnl2 | Hsph1 | Mapk8ip3 | Oxr1 | Ptprz1 | Slc8a1 | Twf1 | |
| Arhgap21 | Camsap2 | Ddx3y | Fry | Htt | Mapre3 | Pabpc1 | Pura | Slc8a2 | Uba1 | |
| Arhgap23 | Camsap3 | Dgkb | Fscn1 | Huwe1 | Mapt | Pacs1 | Pygb | Snap25 | Ube2o | |
| Arhgap26 | Cand1 | Dgkg | Gabbr1 | Icam5 | Mark1 | Pacsin1 | Rab11fip5 | Snap47 | Ubr4 | |
| Arhgap32 | Canx | Dlat | Gabbr2 | Idh3a | Mark2 | Pafah1b1 | Rab1a | Snap91 | Uhrf1bp1l | |
| Arhgap33 | Cap1 | Dld | Gabra1 | Idh3g | Mark3 | Palm | Rab1b | Snd1 | Unc13a | |
| Arhgap39 | Cap2 | Dlg1 | Gabrb2 | Igsf8 | Mark4 | Palm2 | Rab35 | Snph | Unc80 | |
| Arhgap44 | Capn5 | Dlg2 | Gabrg2 | Ildr2 | Mast1 | Pc | Rab3c | Soga3 | Uqcrc1 | |
| Arhgef12 | Capza1 | Dlg3 | Gad1 | Immt | Mast3 | Pcca | Rac1 | Sorbs1 | Uqcrc2 | |
| Arhgef17 | Capza2 | Dlg4 | Gad2 | Ina | Mbp | Pccb | Ralgapa1 | Sorbs2 | Uqcrfs1 | |

**Supplementary Table 3. Postsynaptic proteomes from mouse models of ASD are enriched in synaptic components and pathways in STRING.**

Pathway IDs and false discovery rates (FDR) for the top 25 most enriched Cellular Components, Reactome Pathways, and Biological Processes from STRING for the postsynaptic proteomes in Experiments 1 and 2.

| EXPERIMENT 1 | | | | EXPERIMENT 2 | | | |
|---|---|---|---|---|---|---|---|
| **Pathway ID** | **Cellular Component** | **FDR** | **Count** | **Pathway ID** | **Cellular Component** | **FDR** | **Count** |
| 97458 | neuron part | 1.87E-174 | *369* | 97458 | neuron part | 2.46E-170 | *329* |
| 45202 | synapse | 8.21E-157 | *282* | 120025 | plasma membrane bounded cell projection | 1.17E-144 | *330* |
| 43005 | neuron projection | 9.65E-150 | *317* | 45202 | synapse | 3.04E-144 | *245* |
| 120025 | plasma membrane bounded cell projection | 3.57E-149 | *373* | 42995 | cell projection | 3.59E-144 | *333* |
| 42995 | cell projection | 8.79E-149 | *377* | 43005 | neuron projection | 3.04E-143 | *280* |
| 44456 | synapse part | 3.77E-146 | *254* | 44456 | synapse part | 1.96E-135 | *222* |
| 120038 | plasma membrane bounded cell projection part | 2.35E-129 | *302* | 120038 | plasma membrane bounded cell projection part | 9.21E-120 | *262* |
| 98794 | postsynapse | 1.25E-116 | *190* | 98794 | postsynapse | 8.16E-112 | *170* |
| 5737 | cytoplasm | 1.67E-116 | *669* | 30424 | axon | 2.49E-97 | *175* |
| 36477 | somatodendritic compartment | 3.64E-108 | *231* | 36477 | somatodendritic compartment | 2.49E-97 | *198* |
| 5623 | cell | 2.32E-106 | *757* | 5737 | cytoplasm | 1.34E-92 | *545* |
| 44464 | cell part | 8.40E-106 | *756* | 71944 | cell periphery | 8.72E-92 | *372* |
| 16020 | membrane | 4.63E-100 | *565* | 30425 | dendrite | 1.60E-88 | *164* |
| 30425 | dendrite | 2.17E-98 | *191* | 44464 | cell part | 1.66E-88 | *622* |
| 44444 | cytoplasmic part | 3.30E-94 | *564* | 99572 | postsynaptic specialization | 5.59E-85 | *116* |
| 71944 | cell periphery | 3.30E-94 | *428* | 16020 | membrane | 6.98E-85 | *467* |
| 30424 | axon | 7.00E-89 | *182* | 32279 | asymmetric synapse | 1.36E-84 | *116* |
| 98590 | plasma membrane region | 6.06E-87 | *216* | 14069 | postsynaptic density | 3.13E-84 | *115* |
| 5886 | plasma membrane | 2.12E-86 | *410* | 5886 | plasma membrane | 1.72E-83 | *355* |
| 30054 | cell junction | 1.93E-83 | *205* | 43209 | myelin sheath | 2.14E-80 | *104* |
| 5622 | intracellular | 1.97E-83 | *697* | 98590 | plasma membrane region | 1.04E-77 | *184* |
| 99572 | postsynaptic specialization | 1.96E-82 | *123* | 30054 | cell junction | 1.76E-76 | *177* |
| 32279 | asymmetric synapse | 5.29E-82 | *123* | 5856 | cytoskeleton | 6.39E-76 | *232* |
| 14069 | postsynaptic density | 9.67E-82 | *122* | 33267 | axon part | 3.91E-73 | *127* |
| 44424 | intracellular part | 4.93E-80 | *686* | 44444 | cytoplasmic part | 1.17E-69 | *450* |

| EXPERIMENT 1 | | | | EXPERIMENT 2 | | | |
|---|---|---|---|---|---|---|---|
| **Pathway ID** | **Reactome Pathway** | **FDR** | **Count** | **Pathway ID** | **Reactome Pathway** | **FDR** | **Count** |
| 112316 | Neuronal System | 1.05E-53 | *99* | 112316 | Neuronal System | 1.74E-43 | *80* |
| 112315 | Transmission across Chemical Synapses | 9.10E-36 | *65* | 112315 | Transmission across Chemical Synapses | 2.96E-32 | *56* |
| 6794362 | Protein-protein interactions at synapses | 1.41E-29 | *40* | 6794362 | Protein-protein interactions at synapses | 6.04E-25 | *33* |
| 422475 | Axon guidance | 1.54E-27 | *62* | 194315 | Signaling by Rho GTPases | 2.03E-24 | *61* |
| 1266738 | Developmental Biology | 1.81E-24 | *76* | 422475 | Axon guidance | 6.53E-24 | *52* |
| 112314 | Neurotransmitter receptors and postsynaptic signal transmission | 3.05E-24 | *44* | 195258 | RHO GTPase Effectors | 1.24E-20 | *47* |
| 194315 | Signaling by Rho GTPases | 3.05E-24 | *67* | 373760 | L1CAM interactions | 1.24E-20 | *29* |
| 195258 | RHO GTPase Effectors | 6.38E-24 | *56* | 199991 | Membrane Trafficking | 1.99E-20 | *67* |
| 373760 | L1CAM interactions | 2.36E-23 | *34* | 1266738 | Developmental Biology | 2.52E-20 | *62* |
| 199991 | Membrane Trafficking | 3.58E-21 | *76* | 5653656 | Vesicle-mediated transport | 2.47E-19 | *67* |
| 6794361 | Neurexins and neuroligins | 1.49E-20 | *25* | 6794361 | Neurexins and neuroligins | 6.97E-18 | *21* |
| 1428517 | The citric acid (TCA) cycle and respiratory electron transport | 3.67E-18 | *37* | 112314 | Neurotransmitter receptors and postsynaptic signal transmission | 1.04E-17 | *33* |
| 437239 | Recycling pathway of L1 | 9.83E-18 | *23* | 2132295 | MHC class II antigen presentation | 4.86E-17 | *31* |
| 399719 | Trafficking of AMPA receptors | 3.19E-16 | *20* | 162582 | Signal Transduction | 3.25E-16 | *149* |
| 162582 | Signal Transduction | 3.62E-16 | *172* | 437239 | Recycling pathway of L1 | 3.25E-16 | *20* |
| 2029482 | Regulation of actin dynamics for phagocytic cup formation | 2.97E-14 | *23* | 399719 | Trafficking of AMPA receptors | 1.39E-15 | *18* |
| 5683057 | MAPK family signaling cascades | 5.20E-14 | *42* | 399721 | Glutamate binding, activation of AMPA receptors and synaptic plasticity | 1.39E-15 | *18* |
| 168256 | Immune System | 7.74E-14 | *120* | 3371497 | HSP90 chaperone for steroid hormone receptor | 6.22E-15 | *21* |
| 2029480 | Fcgamma receptor (FCGR) dependent phagocytosis | 1.10E-13 | *25* | 168256 | Immune System | 6.74E-15 | *107* |
| 442755 | Activation of NMDA receptors and postsynaptic events | 1.21E-13 | *19* | 438066 | Unblocking of NMDA receptors, glutamate binding and activation | 6.04E-14 | *15* |
| 3371497 | HSP90 chaperone cycle for steroid hormone receptors (SHR) | 2.81E-13 | *21* | 6807878 | COPI-mediated anterograde transport | 1.08E-13 | *25* |
| 2132295 | MHC class II antigen presentation | 3.01E-13 | *29* | 8856828 | Clathrin-mediated endocytosis | 1.62E-13 | *27* |
| 5663213 | RHO GTPases Activate WASPs and WAVEs | 7.74E-13 | *18* | 442755 | Activation of NMDA receptors and postsynaptic events | 5.65E-12 | *16* |
| 438066 | Unblocking of NMDA receptors, glutamate binding and activation | 9.96E-13 | *15* | 442982 | Ras activation upon Ca2+ influx through NMDA receptor | 1.38E-11 | *13* |
| 168249 | Innate Immune System | 2.21E-12 | *81* | 8849932 | Synaptic adhesion-like molecules | 5.87E-11 | *13* |

| EXPERIMENT 1 | | | | EXPERIMENT 2 | | | |
|---|---|---|---|---|---|---|---|
| **Pathway ID** | **Biological Process** | **FDR** | **Count** | **Pathway ID** | **Biological Process** | **FDR** | **Count** |
| 9987 | cellular process | 2.98E-89 | *707* | 16043 | cellular component organization | 7.92E-84 | *368* |
| 71840 | cellular component organization or biogenesis | 2.38E-81 | *424* | 71840 | cellular component organization or biogenesis | 2.54E-80 | *369* |
| 16043 | cellular component organization | 1.63E-78 | *411* | 9987 | cellular process | 3.20E-73 | *580* |
| 65008 | regulation of biological quality | 5.10E-75 | *348* | 7399 | nervous system development | 2.06E-70 | *240* |
| 7399 | nervous system development | 4.03E-71 | *271* | 48699 | generation of neurons | 3.49E-63 | *194* |
| 22008 | neurogenesis | 1.42E-61 | *222* | 51128 | regulation of cellular component organization | 4.05E-63 | *237* |
| 48699 | generation of neurons | 2.40E-61 | *214* | 22008 | neurogenesis | 5.56E-63 | *200* |
| 51179 | localization | 1.65E-58 | *363* | 7010 | cytoskeleton organization | 1.19E-58 | *148* |
| 51128 | regulation of cellular component organization | 4.65E-57 | *257* | 51179 | localization | 1.30E-57 | *315* |
| 32501 | multicellular organismal process | 6.30E-54 | *424* | 65008 | regulation of biological quality | 2.00E-56 | *276* |
| 48731 | system development | 2.89E-52 | *339* | 32879 | regulation of localization | 5.11E-54 | *234* |
| 65007 | biological regulation | 4.38E-52 | *580* | 32501 | multicellular organismal process | 3.59E-53 | *365* |
| 50804 | modulation of chemical synaptic transmission | 1.51E-51 | *104* | 51641 | cellular localization | 3.66E-53 | *193* |
| 32879 | regulation of localization | 3.21E-51 | *260* | 120035 | regulation of plasma membrane bounded cell projection organization | 2.27E-52 | *125* |
| 7010 | cytoskeleton organization | 4.44E-51 | *153* | 65007 | biological regulation | 2.24E-51 | *492* |
| 51049 | regulation of transport | 7.94E-49 | *209* | 48731 | system development | 6.68E-50 | *291* |
| 48856 | anatomical structure development | 1.27E-48 | *370* | 48856 | anatomical structure development | 3.74E-48 | *320* |
| 51234 | establishment of localization | 1.29E-48 | *294* | 31175 | neuron projection development | 1.97E-47 | *113* |
| 51641 | cellular localization | 1.54E-48 | *209* | 120036 | plasma membrane bounded cell projection organization | 1.01E-46 | *136* |
| 7275 | multicellular organism development | 3.30E-48 | *355* | 7275 | multicellular organism development | 1.78E-46 | *305* |
| 6810 | transport | 5.48E-47 | *284* | 30030 | cell projection organization | 1.86E-46 | *138* |
| 120035 | regulation of plasma membrane bounded cell projection organization | 1.28E-46 | *130* | 48666 | neuron development | 2.81E-46 | *122* |
| 50789 | regulation of biological process | 2.18E-45 | *546* | 50794 | regulation of cellular process | 2.81E-46 | *451* |
| 32502 | developmental process | 2.21E-45 | *376* | 51234 | establishment of localization | 8.55E-46 | *252* |
| 50794 | regulation of cellular process | 1.26E-44 | *525* | 32502 | developmental process | 1.00E-45 | *326* |

**Supplementary Table 4. Top Diseases and Functions enriched in postsynaptic proteomes from ASD mouse models.**

Top 100 most enriched annotations, with overlap *p*-values, generated in IPA for the postsynaptic proteomes in Experiment 1 and Experiment 2.

| EXPERIMENT 1 | | |
|---|---|---|
| **Diseases or Functions Annotation** | **Overlap *p*-value** | **Molecules #** |
| Organization of cytoplasm | 2.00E-98 | *306* |
| Organization of cytoskeleton | 2.60E-98 | *291* |
| Development of neurons | 1.31E-96 | *219* |
| Neurotransmission | 1.09E-91 | *153* |
| Microtubule dynamics | 5.78E-91 | *260* |
| Neuritogenesis | 9.40E-87 | *182* |
| Morphogenesis of neurons | 9.88E-87 | *183* |
| Synaptic transmission | 1.58E-86 | *135* |
| Formation of cellular protrusions | 2.55E-77 | *212* |
| Motor dysfunction or movement disorder | 8.55E-75 | *227* |
| Movement Disorders | 2.04E-72 | *222* |
| Morphology of neurons | 3.98E-70 | *162* |
| Morphology of nervous system | 2.90E-65 | *199* |
| Familial encephalopathy | 3.33E-60 | *235* |
| Potentiation of synapse | 1.09E-55 | *98* |
| Cognition | 1.11E-55 | *127* |
| Learning | 2.04E-55 | *121* |
| Long-term potentiation | 5.40E-55 | *97* |
| Branching of neurons | 3.22E-54 | *103* |
| Cognitive impairment | 1.35E-53 | *149* |
| Disorder of basal ganglia | 5.25E-53 | *156* |
| Neuromuscular disease | 7.91E-53 | *172* |
| Branching of neurites | 1.34E-51 | *99* |
| Cell-cell contact | 4.80E-50 | *152* |
| Neurological signs | 5.80E-49 | *144* |
| Abnormal morphology of neurons | 3.13E-48 | *120* |
| Dendritic growth/branching | 3.07E-47 | *85* |
| Branching of cells | 6.15E-47 | *106* |
| Synaptic depression | 2.87E-46 | *62* |
| Organization of cellular membrane | 2.99E-46 | *84* |
| Abnormal morphology of nervous system | 3.20E-46 | *154* |
| Sprouting | 3.56E-45 | *107* |
| Thyroid carcinoma | 7.30E-45 | *623* |
| Growth of neurites | 8.93E-45 | *110* |
| Endocrine carcinoma | 1.31E-44 | *624* |
| Neck neoplasm | 1.44E-44 | *628* |
| Thyroid gland tumor | 2.02E-44 | *624* |
| Formation of cell-cell contacts | 2.61E-44 | *85* |
| Nonpituitary endocrine tumor | 3.90E-44 | *627* |
| Proliferation of neuronal cells | 5.18E-44 | *119* |
| Formation of intercellular junctions | 6.41E-44 | *84* |
| Cancer of secretory structure | 1.87E-43 | *660* |
| Head and neck tumor | 2.88E-43 | *646* |
| Head and neck cancer | 7.61E-43 | *641* |
| Assembly of intercellular junctions | 1.84E-42 | *80* |
| Head and neck carcinoma | 2.19E-42 | *638* |
| Formation of plasma membrane | 4.24E-42 | *81* |
| Dyskinesia | 4.76E-42 | *122* |
| Seizures | 6.27E-42 | *108* |
| Excitatory postsynaptic potential | 7.23E-42 | *55* |
| Progressive neurological disorder | 1.03E-41 | *174* |
| Non-melanoma solid tumor | 2.15E-41 | *727* |
| Non-hematological solid tumor | 2.37E-41 | *731* |
| Tumorigenesis of tissue | 2.87E-41 | *725* |
| Plasticity of synapse | 1.80E-40 | *55* |
| Conditioning | 2.39E-40 | *70* |
| Nonhematologic malignant neoplasm | 2.54E-40 | *728* |
| Carcinoma | 8.90E-40 | *721* |
| Seizure disorder | 1.36E-39 | *112* |
| Cancer | 2.25E-39 | *737* |
| Development of gap junctions | 5.08E-39 | *72* |
| Malignant solid tumor | 1.80E-38 | *734* |
| Synaptic transmission of cells | 2.12E-38 | *53* |
| Solid tumor | 2.76E-38 | *736* |
| Developmental process of synapse | 4.27E-38 | *69* |
| Transport of molecule | 6.19E-38 | *218* |
| Huntington Disease | 1.47E-37 | *110* |
| Progressive encephalopathy | 4.01E-36 | *150* |
| Abdominal neoplasm | 7.29E-36 | *693* |
| Extracranial solid tumor | 8.95E-36 | *730* |
| Morphology of neurites | 3.02E-35 | *69* |
| Intestinal adenocarcinoma | 3.51E-35 | *583* |
| Autism spectrum disorder or intellectual disability | 6.74E-35 | *110* |
| Gastrointestinal adenocarcinoma | 7.66E-35 | *592* |
| Large intestine adenocarcinoma | 8.15E-35 | *582* |
| Abdominal carcinoma | 1.08E-34 | *671* |
| Intestinal carcinoma | 1.20E-34 | *585* |
| Large intestine carcinoma | 2.69E-34 | *584* |
| Organization of organelle | 4.24E-34 | *116* |
| Morphology of cellular protrusions | 6.67E-34 | *77* |
| Severe psychological disorder | 1.71E-33 | *118* |
| Abdominal cancer | 2.67E-33 | *683* |
| Genitourinary tumor | 2.84E-33 | *543* |
| Outgrowth of neurites | 4.99E-33 | *85* |
| Adenocarcinoma | 6.35E-33 | *650* |
| Digestive organ tumor | 8.12E-33 | *664* |
| Anogenital cancer | 1.19E-32 | *506* |
| Abdominal adenocarcinoma | 1.28E-32 | *640* |
| Axonogenesis | 1.35E-32 | *66* |
| Action potential of cells | 1.82E-32 | *52* |
| Action potential of neurons | 3.47E-32 | *49* |
| Malignant genitourinary solid tumor | 3.81E-32 | *533* |
| Outgrowth of cells | 1.03E-31 | *88* |
| Malignant neoplasm of large intestine | 2.03E-31 | *597* |
| Quantity of nervous tissue | 2.42E-31 | *90* |
| Gastrointestinal carcinoma | 2.47E-31 | *598* |
| Large intestine neoplasm | 3.88E-31 | *598* |
| Dementia | 4.55E-31 | *110* |
| Schizophrenia spectrum disorder | 6.47E-31 | *93* |
| Schizophrenia | 7.11E-31 | *90* |

| EXPERIMENT 2 | | |
|---|---|---|
| **Diseases or Functions Annotation** | **Overlap *p*-value** | **Molecules #** |
| Organization of cytoskeleton | 2.13E-115 | *324* |
| Organization of cytoplasm | 1.23E-113 | *338* |
| Microtubule dynamics | 1.83E-104 | *287* |
| Development of neurons | 1.88E-101 | *232* |
| Morphogenesis of neurons | 4.79E-95 | *198* |
| Neuritogenesis | 3.33E-94 | *196* |
| Formation of cellular protrusions | 1.49E-88 | *234* |
| Morphology of neurons | 2.40E-78 | *177* |
| Neurotransmission | 7.34E-77 | *143* |
| Synaptic transmission | 5.40E-76 | *129* |
| Morphology of nervous system | 2.29E-71 | *215* |
| Motor dysfunction or movement disorder | 2.32E-66 | *224* |
| Movement Disorders | 7.07E-65 | *220* |
| Thyroid carcinoma | 8.16E-65 | *697* |
| Thyroid gland tumor | 2.92E-64 | *698* |
| Neck neoplasm | 8.89E-64 | *701* |
| Nonpituitary endocrine tumor | 1.26E-62 | *699* |
| Branching of neurons | 7.41E-62 | *114* |
| Familial encephalopathy | 1.70E-61 | *247* |
| Branching of neurites | 2.87E-60 | *111* |
| Shape change of neurites | 5.47E-60 | *112* |
| Head and neck carcinoma | 9.05E-60 | *709* |
| Head and neck tumor | 7.03E-59 | *714* |
| Gastrointestinal adenocarcinoma | 2.39E-55 | *671* |
| Intestinal adenocarcinoma | 8.95E-55 | *660* |
| Dendritic growth/branching | 9.14E-55 | *95* |
| Intestinal carcinoma | 1.31E-54 | *663* |
| Cancer of secretory structure | 1.63E-54 | *721* |
| Large intestine adenocarcinoma | 2.58E-54 | *659* |
| Large intestine carcinoma | 3.66E-54 | *662* |
| Branching of cells | 2.23E-53 | *117* |
| Abnormal morphology of neurons | 6.22E-53 | *130* |
| Sprouting | 2.57E-51 | *118* |
| Abdominal adenocarcinoma | 1.24E-50 | *716* |
| Gastrointestinal carcinoma | 3.48E-50 | *676* |
| Learning | 1.58E-49 | *118* |
| Adenocarcinoma | 1.14E-48 | *722* |
| Tumorigenesis of epithelial neoplasm | 3.06E-48 | *597* |
| Abnormal morphology of nervous system | 3.32E-48 | *163* |
| Malignant neoplasm of large intestine | 3.48E-48 | *671* |
| Cell-cell contact | 4.03E-48 | *155* |
| Cognition | 4.13E-48 | *122* |
| Synaptic depression | 4.38E-48 | *65* |
| Development of carcinoma | 5.52E-48 | *591* |
| Development of malignant tumor | 7.87E-48 | *596* |
| Large intestine neoplasm | 8.13E-48 | *672* |
| Proliferation of neuronal cells | 1.33E-47 | *128* |
| Anogenital cancer | 1.35E-47 | *572* |
| Frequency of tumor | 1.56E-47 | *602* |
| Malignant genitourinary solid tumor | 3.00E-47 | *601* |
| Abdominal carcinoma | 3.61E-47 | *737* |
| Transport of molecule | 1.36E-46 | *245* |
| Genitourinary tumor | 1.55E-46 | *607* |
| Organization of cellular membrane | 8.61E-46 | *86* |
| Growth of neurites | 9.48E-46 | *115* |
| Incidence of tumor | 1.26E-45 | *610* |
| Disorder of basal ganglia | 1.99E-45 | *151* |
| Neuromuscular disease | 4.23E-45 | *167* |
| Gastrointestinal tumor | 6.23E-45 | *686* |
| Gastrointestinal tract cancer | 6.81E-45 | *684* |
| Genitourinary adenocarcinoma | 5.96E-44 | *501* |
| Morphology of cellular protrusions | 8.98E-44 | *91* |
| Seizures | 1.21E-43 | *114* |
| Pelvic adenocarcinoma | 2.57E-43 | *468* |
| Cognitive impairment | 2.91E-43 | *140* |
| Potentiation of synapse | 1.76E-42 | *87* |
| Genital tumor | 2.40E-42 | *524* |
| Digestive system cancer | 4.85E-42 | *717* |
| Abdominal cancer | 8.93E-42 | *743* |
| Pelvic tumor | 1.17E-41 | *547* |
| Genital tract cancer | 1.31E-41 | *516* |
| Pelvic cancer | 3.41E-41 | *540* |
| Neurological signs | 4.34E-41 | *138* |
| Seizure disorder | 5.48E-41 | *118* |
| Long-term potentiation | 6.08E-41 | *85* |
| Morphology of neurites | 9.75E-41 | *77* |
| Breast or pancreatic cancer | 1.53E-40 | *435* |
| Digestive organ tumor | 1.92E-40 | *722* |
| Progressive neurological disorder | 2.47E-40 | *179* |
| Abdominal neoplasm | 3.01E-40 | *746* |
| Development of digestive organ tumor | 5.23E-40 | *446* |
| Development of adenocarcinoma | 5.37E-40 | *427* |
| Breast or gynecological cancer | 6.42E-40 | *466* |
| Genitourinary carcinoma | 1.92E-39 | *532* |
| Formation of cell-cell contacts | 1.86E-37 | *80* |
| Progressive encephalopathy | 4.27E-37 | *158* |
| Colon carcinoma | 5.67E-37 | *387* |
| Breast or colorectal cancer | 2.87E-36 | *498* |
| Formation of intercellular junctions | 3.26E-36 | *78* |
| Assembly of intercellular junctions | 1.05E-35 | *75* |
| Severe psychological disorder | 1.33E-35 | *126* |
| Carcinoma | 1.70E-35 | *762* |
| Colon tumor | 1.72E-35 | *400* |
| Hepatobiliary carcinoma | 1.73E-35 | *406* |
| Development of genital tumor | 2.15E-35 | *381* |
| Formation of plasma membrane | 2.17E-35 | *76* |
| Development of colorectal tumor | 3.58E-35 | *397* |
| Liver tumor | 3.66E-35 | *443* |
| Colorectal carcinoma | 7.32E-35 | *396* |
| Colon cancer | 7.36E-35 | *395* |

**Supplementary Table 5. Summary of Upstream Regulators, Downstream Effects, and Canonical Pathways enriched in postsynaptic proteomes from ASD mouse models.**

Enrichment *p*-values for the top 5 most enriched Upstream Regulators, Molecular and Cellular Functions, Physiological Systems, Diseases and Disorders, and Canonical Pathways were generated in IPA for the postsynaptic proteomes in Experiment 1 and Experiment 2.

| EXPERIMENT 1 | | |
|---|---|---|
| **Top Upstream Regulators** | **Overlap *p*-value** | **Regulators in Mechanistic Network** |
| **HTT** | 8.31E-42 | |
| **MLXIPL** | 1.65E-27 | |
| **MYCN** | 1.16E-26 | |
| **MYC** | 2.21E-24 | CTNNB1, MYC |
| **HDAC4** | 1.33E-23 | |

| EXPERIMENT 2 | | |
|---|---|---|
| **Top Upstream Regulators** | **Overlap *p*-value** | **Regulators in Mechanistic Network** |
| **HTT** | 1.85E-37 | |
| **MMP12** | 7.76E-21 | |
| **TP53** | 2.43E-20 | |
| **SMYD1** | 4.98E-17 | |
| **HDAC4** | 6.22E-17 | |

| EXPERIMENT 1 | | |
|---|---|---|
| **Top Molecular and Cellular Functions** | ***p*-value range** | **Number of Molecules** |
| **Cellular Assembly and Organization** | 2.69E-11 - 2.00E-98 | 422 |
| **Cellular Function and Maintenance** | 2.69E-11 - 2.00E-98 | 460 |
| **Cellular Development** | 6.01E-13 - 1.31E-96 | 247 |
| **Cellular Growth and Proliferation** | 6.01E-13 - 1.31E-96 | 247 |
| **Cell-To-Cell Signaling and Interaction** | 1.38E-11 - 1.09E-91 | 306 |

| EXPERIMENT 2 | | |
|---|---|---|
| **Top Molecular and Cellular Functions** | ***p*-value range** | **Number of Molecules** |
| **Cellular Assembly and Organization** | 1.34E-11 - 2.13E-115 | 451 |
| **Cellular Function and Maintenance** | 8.35E-12 - 2.13E-115 | 499 |
| **Cellular Development** | 7.70E-12 - 1.88E-101 | 291 |
| **Cellular Growth and Proliferation** | 5.85E-16 - 1.88E-101 | 269 |
| **Cell Morphology** | 1.50E-11 - 4.79E-95 | 361 |

| EXPERIMENT 1 | | |
|---|---|---|
| **Top Physiological Functions** | ***p*-value range** | **Number of Molecules** |
| **Nervous System Development and Function** | 1.87E-11 - 1.31E-96 | 418 |
| **Tissue Development** | 2.37E-12 - 1.31E-96 | 314 |
| **Organismal Development** | 1.87E-11 - 9.40E-87 | 292 |
| **Tissue Morphology** | 7.55E-12 - 3.98E-70 | 266 |
| **Behavior** | 3.77E-12 - 1.11E-55 | 208 |

| EXPERIMENT 2 | | |
|---|---|---|
| **Top Physiological Functions** | ***p*-value range** | **Number of Molecules** |
| **Nervous System Development and Function** | 1.50E-11 - 1.88E-101 | 435 |
| **Tissue Development** | 1.09E-11 - 1.88E-101 | 340 |
| **Organismal Development** | 1.09E-11 - 4.79E-95 | 314 |
| **Tissue Morphology** | 1.50E-11 - 2.40E-78 | 225 |
| **Embryonic Development** | 1.09E-11 - 2.87E-60 | 150 |

| EXPERIMENT 1 | | |
|---|---|---|
| **Top Diseases and Disorders** | ***p*-value range** | **Number of Molecules** |
| **Neurological Disease** | 2.43E-11 - 8.55E-75 | 502 |
| **Hereditary Disorder** | 1.71E-11 - 3.33E-60 | 269 |
| **Organismal Injury and Abnormalities** | 2.49E-11 - 3.33E-60 | 751 |
| **Psychological Disorders** | 2.40E-12 - 5.25E-53 | 304 |
| **Skeletal and Muscular Disorders** | 1.71E-11 - 7.91E-53 | 242 |

| EXPERIMENT 2 | | |
|---|---|---|
| **Top Diseases and Disorders** | ***p*-value range** | **Number of Molecules** |
| **Neurological Disease** | 1.53E-11 - 2.32E-66 | 526 |
| **Cancer** | 1.53E-11 - 8.16E-65 | 771 |
| **Endocrine System Disorders** | 1.97E-15 - 8.16E-65 | 723 |
| **Organismal Injury and Abnormalities** | 1.53E-11 - 8.16E-65 | 780 |
| **Hereditary Disorder** | 7.16E-13 - 1.70E-61 | 285 |

| EXPERIMENT 1 | | |
|---|---|---|
| **Top Canonical Pathways** | **Overlap *p*-value** | **Ratio (%)** |
| **Synaptogenesis Signaling** | 2.82E-57 | 89/312 (28.5%) |
| **Remodeling of Epithelial Adherens Junctions** | 1.13E-30 | 33/68 (48.5%) |
| **Opioid Signaling** | 1.77E-30 | 56/247 (22.7%) |
| **CREB Signaling in Neurons** | 1.06E-28 | 50/207 (24.2%) |
| **Endocannabinoid Neuronal Synapse** | 9.32E-27 | 39/128 (30.5%) |

| EXPERIMENT 2 | | |
|---|---|---|
| **Top Canonical Pathways** | **Overlap *p*-value** | **Ratio** |
| **Synaptogenesis Signaling** | 1.76E-51 | 86/312 (27.6%) |
| **Remodeling of Epithelial Adherens Junctions** | 1.49E-25 | 30/68 (44.1%) |
| **Opioid Signaling** | 1.92E-24 | 51/247 (20.6%) |
| **CREB Signaling in Neurons** | 1.42E-23 | 46/207 (22.2%) |
| **Endocannabinoid Neuronal Synapse** | 1.92E-22 | 36/128 (28.1%) |

**Supplementary Table 6. Statistical analysis for Figure 8.**

All statistical analyses were performed in JMP version 14 (SAS). SEM = standard error of mean, CL = confidence limit, Dif = difference, DF = degrees of freedom.

| | | | | | | | | | | | | | | |
|---|---|---|---|---|---|---|---|---|---|---|---|---|---|---|
| **Figure 8B PSD Fraction** | **Rac1 (% Control)** | **Fmr1 KO** | | | | | | | | | **Difference** | 67.04 | t Ratio | 3.0183 |
| | | | **Level** | **N** | **Mean** | **Std Dev** | **SEM** | **Lower 95%** | **Upper 95%** | ***Student's t-Test*** | **Std Err Dif** | 22.21 | DF | 10.0000 |
| | | | C57BL6J | 6 | 100.00 | 26.88 | 10.97 | 71.79 | 128.21 | | **Upper CL Dif** | 116.53 | Prob > \|t\| | 0.0129* |
| | | | Fmr1 KO | 6 | 167.04 | 47.30 | 19.31 | 117.40 | 216.68 | | **Lower CL Dif** | 17.55 | Prob > t | 0.0065** |
| | | | | | | | | | | | **Confidence** | 0.95 | Prob < t | 0.9935 |
| | | **Pten Het** | | | | | | | | | **Difference** | 18.69 | t Ratio | 2.0741 |
| | | | **Level** | **N** | **Mean** | **Std Dev** | **SEM** | **Lower 95%** | **Upper 95%** | ***Student's t-Test*** | **Std Err Dif** | 9.01 | DF | 10 |
| | | | C57BL6J | 6 | 100.00 | 15.57 | 6.36 | 83.66 | 116.34 | | **Upper CL Dif** | 38.77 | Prob > \|t\| | 0.0648 |
| | | | Pten Het | 6 | 118.69 | 15.64 | 6.39 | 102.27 | 135.11 | | **Lower CL Dif** | -1.39 | Prob > t | 0.0324* |
| | | | | | | | | | | | **Confidence** | 0.95 | Prob < t | 0.9676 |
| | | **Cntnap2 KO** | | | | | | | | | **Difference** | 18.86 | t Ratio | 0.9368 |
| | | | **Level** | **N** | **Mean** | **Std Dev** | **SEM** | **Lower 95%** | **Upper 95%** | ***Student's t-Test*** | **Std Err Dif** | 20.13 | DF | 10 |
| | | | C57BL6J | 6 | 100.00 | 26.88 | 10.97 | 71.79 | 128.21 | | **Upper CL Dif** | 63.71 | Prob > \|t\| | 0.3709 |
| | | | Cntnap2 KO | 6 | 118.86 | 41.34 | 16.88 | 75.48 | 162.24 | | **Lower CL Dif** | -25.99 | Prob > t | 0.1855 |
| | | | | | | | | | | | **Confidence** | 0.95 | Prob < t | 0.8145 |
| | | **BTBR+** | | | | | | | | | **Difference** | -14.61 | t Ratio | -0.7217 |
| | | | **Level** | **N** | **Mean** | **Std Dev** | **SEM** | **Lower 95%** | **Upper 95%** | ***Student's t-Test*** | **Std Err Dif** | 20.24 | DF | 10 |
| | | | C57BL6J | 6 | 100.00 | 26.88 | 10.97 | 71.79 | 128.21 | | **Upper CL Dif** | 30.49 | Prob > \|t\| | 0.4870 |
| | | | BTBR+ | 6 | 85.39 | 41.66 | 17.01 | 41.68 | 129.11 | | **Lower CL Dif** | -59.70 | Prob > t | 0.7565 |
| | | | | | | | | | | | **Confidence** | 0.95 | Prob < t | 0.2435 |
| | | **Cacna1c*** | | | | | | | | | **Difference** | 0.53 | t Ratio | 0.0240 |
| | | | **Level** | **N** | **Mean** | **Std Dev** | **SEM** | **Lower 95%** | **Upper 95%** | ***Student's t-Test*** | **Std Err Dif** | 22.25 | DF | 4 |
| | | | C57BL6J | 3 | 100.00 | 36.12 | 20.85 | 10.27 | 189.73 | | **Upper CL Dif** | 62.30 | Prob > \|t\| | 0.9820 |
| | | | Cacna1c G406R | 3 | 100.53 | 13.42 | 7.75 | 67.20 | 133.86 | | **Lower CL Dif** | -61.23 | Prob > t | 0.4910 |
| | | | | | | | | | | | **Confidence** | 0.95 | Prob < t | 0.5090 |
| | | **Shank3*** | | | | | | | | | **Difference** | 3.67 | t Ratio | 0.2346 |
| | | | **Level** | **N** | **Mean** | **Std Dev** | **SEM** | **Lower 95%** | **Upper 95%** | ***Student's t-Test*** | **Std Err Dif** | 15.65 | DF | 4 |
| | | | Shank3 WT | 3 | 100.00 | 21.06 | 12.16 | 47.69 | 152.31 | | **Upper CL Dif** | 47.13 | Prob > \|t\| | 0.8260 |
| | | | Shank3 InsG3680 | 3 | 103.67 | 17.07 | 9.85 | 61.27 | 146.07 | | **Lower CL Dif** | -39.78 | Prob > t | 0.4130 |
| | | | | | | | | | | | **Confidence** | 0.95 | Prob < t | 0.5870 |
| | | **Anks1b Het** | | | | | | | | | **Difference** | 22.23 | t Ratio | 2.5091 |
| | | | **Level** | **N** | **Mean** | **Std Dev** | **SEM** | **Lower 95%** | **Upper 95%** | ***Student's t-Test*** | **Std Err Dif** | 8.86 | DF | 10 |
| | | | Anks1b WT | 6 | 100.00 | 15.50 | 6.33 | 83.73 | 116.27 | | **Upper CL Dif** | 41.98 | Prob > \|t\| | 0.0310* |
| | | | Anks1b Het | 6 | 122.23 | 15.19 | 6.20 | 106.29 | 138.18 | | **Lower CL Dif** | 2.49 | Prob > t | 0.0155* |
| | | | | | | | | | | | **Confidence** | 0.95 | Prob < t | 0.9845 |
| **Figure 8C Total Lysate** | **Rac1 (% Control)** | **Fmr1 KO** | | | | | | | | | **Difference** | -17.15 | t Ratio | -8.6294 |
| | | | **Level** | **N** | **Mean** | **Std Dev** | **SEM** | **Lower 95%** | **Upper 95%** | ***Student's t-Test*** | **Std Err Dif** | 1.99 | DF | 10 |
| | | | C57BL6J | 6 | 100.00 | 2.09 | 0.85 | 97.80 | 102.20 | | **Upper CL Dif** | -12.72 | Prob > \|t\| | <.0001*** |
| | | | Fmr1 KO | 6 | 82.85 | 4.40 | 1.79 | 78.24 | 87.46 | | **Lower CL Dif** | -21.58 | Prob > t | 1.0000 |
| | | | | | | | | | | | **Confidence** | 0.95 | Prob < t | <.0001*** |
| | | **Pten Het** | | | | | | | | | **Difference** | -27.97 | t Ratio | -6.9700 |
| | | | **Level** | **N** | **Mean** | **Std Dev** | **SEM** | **Lower 95%** | **Upper 95%** | ***Student's t-Test*** | **Std Err Dif** | 4.01 | DF | 12 |
| | | | C57BL6J | 7 | 100.00 | 7.50 | 2.83 | 93.07 | 106.93 | | **Upper CL Dif** | -19.23 | Prob > \|t\| | <.0001*** |
| | | | Pten Het | 7 | 72.03 | 7.52 | 2.84 | 65.08 | 78.98 | | **Lower CL Dif** | -36.71 | Prob > t | 1.0000 |
| | | | | | | | | | | | **Confidence** | 0.95 | Prob < t | <.0001*** |
| | | **Cntnap2 KO** | | | | | | | | | **Difference** | -17.30 | t Ratio | -3.9926 |
| | | | **Level** | **N** | **Mean** | **Std Dev** | **SEM** | **Lower 95%** | **Upper 95%** | ***Student's t-Test*** | **Std Err Dif** | 4.33 | DF | 10 |
| | | | C57BL6J | 6 | 100.00 | 2.09 | 0.85 | 97.80 | 102.20 | | **Upper CL Dif** | -7.65 | Prob > \|t\| | 0.0025** |
| | | | Cntnap2 KO | 6 | 82.70 | 10.40 | 4.25 | 71.78 | 93.62 | | **Lower CL Dif** | -26.95 | Prob > t | 0.9987 |
| | | | | | | | | | | | **Confidence** | 0.95 | Prob < t | 0.0013** |
| | | **BTBR+** | | | | | | | | | **Difference** | -26.25 | t Ratio | -5.8941 |
| | | | **Level** | **N** | **Mean** | **Std Dev** | **SEM** | **Lower 95%** | **Upper 95%** | ***Student's t-Test*** | **Std Err Dif** | 4.45 | DF | 10 |
| | | | C57BL6J | 6 | 100.00 | 2.09 | 0.85 | 97.80 | 102.20 | | **Upper CL Dif** | -16.33 | Prob > \|t\| | 0.0002*** |
| | | | BTBR+ | 6 | 73.75 | 10.71 | 4.37 | 62.51 | 84.98 | | **Lower CL Dif** | -36.18 | Prob > t | 0.9999 |
| | | | | | | | | | | | **Confidence** | 0.95 | Prob < t | <.0001*** |
| | | **Cacna1c*** | | | | | | | | | **Difference** | 21.58 | t Ratio | 7.6495 |
| | | | **Level** | **N** | **Mean** | **Std Dev** | **SEM** | **Lower 95%** | **Upper 95%** | ***Student's t-Test*** | **Std Err Dif** | 2.82 | DF | 4 |
| | | | C57BL6J | 3 | 100.00 | 3.12 | 1.80 | 92.25 | 107.75 | | **Upper CL Dif** | 29.41 | Prob > \|t\| | 0.0016** |
| | | | Cacna1c G406R | 3 | 121.58 | 3.76 | 2.17 | 112.23 | 130.93 | | **Lower CL Dif** | 13.75 | Prob > t | 0.0008*** |
| | | | | | | | | | | | **Confidence** | 0.95 | Prob < t | 0.9992 |
| | | **Shank3*** | | | | | | | | | **Difference** | -25.93 | t Ratio | -7.0203 |
| | | | **Level** | **N** | **Mean** | **Std Dev** | **SEM** | **Lower 95%** | **Upper 95%** | ***Student's t-Test*** | **Std Err Dif** | 3.69 | DF | 4 |
| | | | Shank3 WT | 3 | 100.00 | 5.25 | 3.03 | 86.97 | 113.03 | | **Upper CL Dif** | -15.67 | Prob > \|t\| | 0.0022** |
| | | | Shank3 InsG3680 | 3 | 74.07 | 3.66 | 2.11 | 64.98 | 83.16 | | **Lower CL Dif** | -36.18 | Prob > t | 0.9989 |
| | | | | | | | | | | | **Confidence** | 0.95 | Prob < t | 0.0011** |
| | | **Anks1b Het** | | | | | | | | | **Difference** | -15.39 | t Ratio | -2.2542 |
| | | | **Level** | **N** | **Mean** | **Std Dev** | **SEM** | **Lower 95%** | **Upper 95%** | ***Student's t-Test*** | **Std Err Dif** | 6.83 | DF | 10 |
| | | | Anks1b WT | 6 | 100.00 | 13.71 | 5.60 | 85.61 | 114.39 | | **Upper CL Dif** | -0.18 | Prob > \|t\| | 0.0478* |
| | | | Anks1b Het | 6 | 84.61 | 9.57 | 3.91 | 74.57 | 94.66 | | **Lower CL Dif** | -30.59 | Prob > t | 0.9761 |
| | | | | | | | | | | | **Confidence** | 0.95 | Prob < t | 0.0239* |
| **Figure 8D Total Lysate** | **RhoA (% Control)** | **Fmr1 KO** | | | | | | | | | **Difference** | -35.07 | t Ratio | -2.4276 |
| | | | **Level** | **N** | **Mean** | **Std Dev** | **SEM** | **Lower 95%** | **Upper 95%** | ***Student's t-Test*** | **Std Err Dif** | 14.45 | DF | 12 |
| | | | C57BL6J | 8 | 100.04 | 22.30 | 7.88 | 81.40 | 118.68 | | **Upper CL Dif** | -3.59 | Prob > \|t\| | 0.0319* |
| | | | Fmr1 KO | 6 | 64.97 | 31.96 | 13.05 | 31.43 | 98.51 | | **Lower CL Dif** | -66.55 | Prob > t | 0.9841 |
| | | | | | | | | | | | **Confidence** | 0.95 | Prob < t | 0.0159* |
| | | **Pten Het** | | | | | | | | | **Difference** | -28.00 | t Ratio | -3.5794 |
| | | | **Level** | **N** | **Mean** | **Std Dev** | **SEM** | **Lower 95%** | **Upper 95%** | ***Student's t-Test*** | **Std Err Dif** | 7.82 | DF | 12 |
| | | | C57BL6J | 7 | 104.28 | 5.37 | 2.03 | 99.31 | 109.25 | | **Upper CL Dif** | -10.96 | Prob > \|t\| | 0.0038* |
| | | | Pten Het | 7 | 76.27 | 19.99 | 7.55 | 57.79 | 94.76 | | **Lower CL Dif** | -45.05 | Prob > t | 0.9981 |
| | | | | | | | | | | | **Confidence** | 0.95 | Prob < t | 0.0019* |
| | | **Cntnap2 KO** | | | | | | | | | **Difference** | -4.24 | t Ratio | -0.3119 |
| | | | **Level** | **N** | **Mean** | **Std Dev** | **SEM** | **Lower 95%** | **Upper 95%** | ***Student's t-Test*** | **Std Err Dif** | 13.61 | DF | 12 |
| | | | C57BL6J | 8 | 100.04 | 22.30 | 7.88 | 81.40 | 118.68 | | **Upper CL Dif** | 25.40 | Prob > \|t\| | 0.7605 |
| | | | Cntnap2 KO | 6 | 95.80 | 28.76 | 11.74 | 65.61 | 125.98 | | **Lower CL Dif** | -33.89 | Prob > t | 0.6198 |
| | | | | | | | | | | | **Confidence** | 0.95 | Prob < t | 0.3802 |
| | | **BTBR+** | | | | | | | | | **Difference** | -14.91 | t Ratio | -1.0506 |
| | | | **Level** | **N** | **Mean** | **Std Dev** | **SEM** | **Lower 95%** | **Upper 95%** | ***Student's t-Test*** | **Std Err Dif** | 14.19 | DF | 12 |
| | | | C57BL6J | 8 | 100.04 | 22.30 | 7.88 | 81.40 | 118.68 | | **Upper CL Dif** | 16.01 | Prob > \|t\| | 0.3141 |
| | | | BTBR+ | 6 | 85.13 | 30.99 | 12.65 | 52.61 | 117.66 | | **Lower CL Dif** | -45.82 | Prob > t | 0.8429 |
| | | | | | | | | | | | **Confidence** | 0.95 | Prob < t | 0.1571 |
| | | **Cacna1c*** | | | | | | | | | **Difference** | -22.28 | t Ratio | -1.2636 |
| | | | **Level** | **N** | **Mean** | **Std Dev** | **SEM** | **Lower 95%** | **Upper 95%** | ***Student's t-Test*** | **Std Err Dif** | 17.63 | DF | 6 |
| | | | C57BL6J | 5 | 100.00 | 29.18 | 13.05 | 63.77 | 136.23 | | **Upper CL Dif** | 20.87 | Prob > \|t\| | 0.2532 |
| | | | Cacna1c G406R | 3 | 77.72 | 6.78 | 3.92 | 60.87 | 94.57 | | **Lower CL Dif** | -65.43 | Prob > t | 0.8734 |
| | | | | | | | | | | | **Confidence** | 0.95 | Prob < t | 0.1266 |
| | | **Shank3*** | | | | | | | | | **Difference** | -21.85 | t Ratio | -2.9048 |
| | | | **Level** | **N** | **Mean** | **Std Dev** | **SEM** | **Lower 95%** | **Upper 95%** | ***Student's t-Test*** | **Std Err Dif** | 7.52 | DF | 4 |
| | | | Shank3 WT | 3 | 100.00 | 8.97 | 5.18 | 77.71 | 122.29 | | **Upper CL Dif** | -0.97 | Prob > \|t\| | 0.0439* |
| | | | Shank3 InsG3680 | 3 | 78.15 | 9.44 | 5.45 | 54.70 | 101.61 | | **Lower CL Dif** | -42.73 | Prob > t | 0.9780 |
| | | | | | | | | | | | **Confidence** | 0.95 | Prob < t | 0.0220* |
| | | **Anks1b Het** | | | | | | | | | **Difference** | -11.91 | t Ratio | -0.7979 |
| | | | **Level** | **N** | **Mean** | **Std Dev** | **SEM** | **Lower 95%** | **Upper 95%** | ***Student's t-Test*** | **Std Err Dif** | 14.93 | DF | 10 |
| | | | Anks1b WT | 6 | 100.00 | 31.43 | 12.83 | 67.02 | 132.98 | | **Upper CL Dif** | 21.35 | Prob > \|t\| | 0.4435 |
| | | | Anks1b Het | 6 | 88.09 | 18.70 | 7.63 | 68.46 | 107.71 | | **Lower CL Dif** | -45.18 | Prob > t | 0.7783 |
| | | | | | | | | | | | **Confidence** | 0.95 | Prob < t | 0.2217 |

| Figure | Measure | Genotype | Level | N | Mean | Std Dev | SEM | Lower 95% | Upper 95% | Test | | | | |
|---|---|---|---|---|---|---|---|---|---|---|---|---|---|---|
| Figure 8E Total Lysate | Cdc42 (% Control) | Fmr1 KO | | | | | | | | *Student's t-Test* | **Difference** | -30.38 | t Ratio | -2.3326 |
| | | | **Level** | **N** | **Mean** | **Std Dev** | **SEM** | **Lower 95%** | **Upper 95%** | | **Std Err Dif** | 13.02 | DF | 10 |
| | | | C57BL6J | 6 | 100.00 | 19.02 | 7.76 | 80.04 | 119.96 | | **Upper CL Dif** | -1.36 | Prob > \|t\| | 0.0419* |
| | | | Fmr1 KO | 6 | 69.62 | 25.61 | 10.46 | 42.75 | 96.50 | | **Lower CL Dif** | -59.40 | Prob > t | 0.9791 |
| | | | | | | | | | | | **Confidence** | 0.95 | Prob < t | 0.0209* |
| | | Pten Het | | | | | | | | *Student's t-Test* | **Difference** | -25.67 | t Ratio | -2.4282 |
| | | | **Level** | **N** | **Mean** | **Std Dev** | **SEM** | **Lower 95%** | **Upper 95%** | | **Std Err Dif** | 10.57 | DF | 15 |
| | | | C57BL6J | 10 | 100.00 | 22.04 | 6.97 | 84.23 | 115.77 | | **Upper CL Dif** | -3.14 | Prob > \|t\| | 0.0282* |
| | | | Pten Het | 7 | 74.33 | 20.54 | 7.77 | 55.33 | 93.33 | | **Lower CL Dif** | -48.21 | Prob > t | 0.9859 |
| | | | | | | | | | | | **Confidence** | 0.95 | Prob < t | 0.0141* |
| | | Cntnap2 KO | | | | | | | | *Student's t-Test* | **Difference** | -55.19 | t Ratio | -6.2294 |
| | | | **Level** | **N** | **Mean** | **Std Dev** | **SEM** | **Lower 95%** | **Upper 95%** | | **Std Err Dif** | 8.86 | DF | 10 |
| | | | C57BL6J | 6 | 100.00 | 19.02 | 7.76 | 80.04 | 119.96 | | **Upper CL Dif** | -35.45 | Prob > \|t\| | <.0001*** |
| | | | Cntnap2 KO | 6 | 44.82 | 10.45 | 4.27 | 33.85 | 55.78 | | **Lower CL Dif** | -74.92 | Prob > t | 1.0000 |
| | | | | | | | | | | | **Confidence** | 0.95 | Prob < t | <.0001*** |
| | | BTBR+ | | | | | | | | *Student's t-Test* | **Difference** | -68.42 | t Ratio | -7.1969 |
| | | | **Level** | **N** | **Mean** | **Std Dev** | **SEM** | **Lower 95%** | **Upper 95%** | | **Std Err Dif** | 9.51 | DF | 10 |
| | | | C57BL6J | 6 | 100.00 | 19.02 | 7.76 | 80.04 | 119.96 | | **Upper CL Dif** | -47.23 | Prob > \|t\| | <.0001*** |
| | | | BTBR+ | 6 | 31.58 | 13.44 | 5.49 | 17.48 | 45.68 | | **Lower CL Dif** | -89.60 | Prob > t | 1.0000 |
| | | | | | | | | | | | **Confidence** | 0.95 | Prob < t | <.0001*** |
| | | Cacna1c* | | | | | | | | *Student's t-Test* | **Difference** | -60.52 | t Ratio | -3.7705 |
| | | | **Level** | **N** | **Mean** | **Std Dev** | **SEM** | **Lower 95%** | **Upper 95%** | | **Std Err Dif** | 16.05 | DF | 4 |
| | | | C57BL6J | 3 | 100.00 | 26.68 | 15.40 | 33.72 | 166.28 | | **Upper CL Dif** | -15.96 | Prob > \|t\| | 0.0196* |
| | | | Cacna1c G406R | 3 | 39.48 | 7.80 | 4.51 | 20.10 | 58.87 | | **Lower CL Dif** | -105.08 | Prob > t | 0.9902 |
| | | | | | | | | | | | **Confidence** | 0.95 | Prob < t | 0.0098** |
| | | Shank3* | | | | | | | | *Student's t-Test* | **Difference** | 3.51 | t Ratio | 0.3138 |
| | | | **Level** | **N** | **Mean** | **Std Dev** | **SEM** | **Lower 95%** | **Upper 95%** | | **Std Err Dif** | 11.18 | DF | 4 |
| | | | Shank3 WT | 3 | 100.00 | 3.17 | 1.83 | 92.13 | 107.87 | | **Upper CL Dif** | 34.54 | Prob > \|t\| | 0.7694 |
| | | | Shank3 InsG3680 | 3 | 103.51 | 19.10 | 11.03 | 56.07 | 150.95 | | **Lower CL Dif** | -27.52 | Prob > t | 0.3847 |
| | | | | | | | | | | | **Confidence** | 0.95 | Prob < t | 0.6153 |
| | | Anks1b Het | | | | | | | | *Student's t-Test* | **Difference** | -2.65 | t Ratio | -0.2786 |
| | | | **Level** | **N** | **Mean** | **Std Dev** | **SEM** | **Lower 95%** | **Upper 95%** | | **Std Err Dif** | 9.52 | DF | 10 |
| | | | Anks1b WT | 6 | 100.00 | 22.24 | 9.08 | 76.66 | 123.34 | | **Upper CL Dif** | 18.56 | Prob > \|t\| | 0.7862 |
| | | | Anks1b Het | 6 | 97.35 | 7.02 | 2.87 | 89.98 | 104.71 | | **Lower CL Dif** | -23.87 | Prob > t | 0.6069 |
| | | | | | | | | | | | **Confidence** | 0.95 | Prob < t | 0.3931 |

31

**Supplementary Figure 1. Tandem-mass-tag (TMT) mass spectrometry and peptide identification shows consistent labeling and quantification of proteins among reporter ions.** **A**) Workflow of proteomics experiments from sample preparation to peptide identification. **B**) Distribution of protein intensities for the 10 reporter ions in Experiment 1, each showing a normal distribution. **C**) Scatterplots comparing reporter ion intensities for proteins between each of the 10 TMT channels. Spearman's rank-order correlation, $\rho > 0.96$ for all TMT channel comparisons. (**D**) For Experiment 2, reporter ion intensities also show normal distributions, and (**E**) Spearman's rank-order correlation, $\rho > 0.96$ for all TMT channel comparisons.

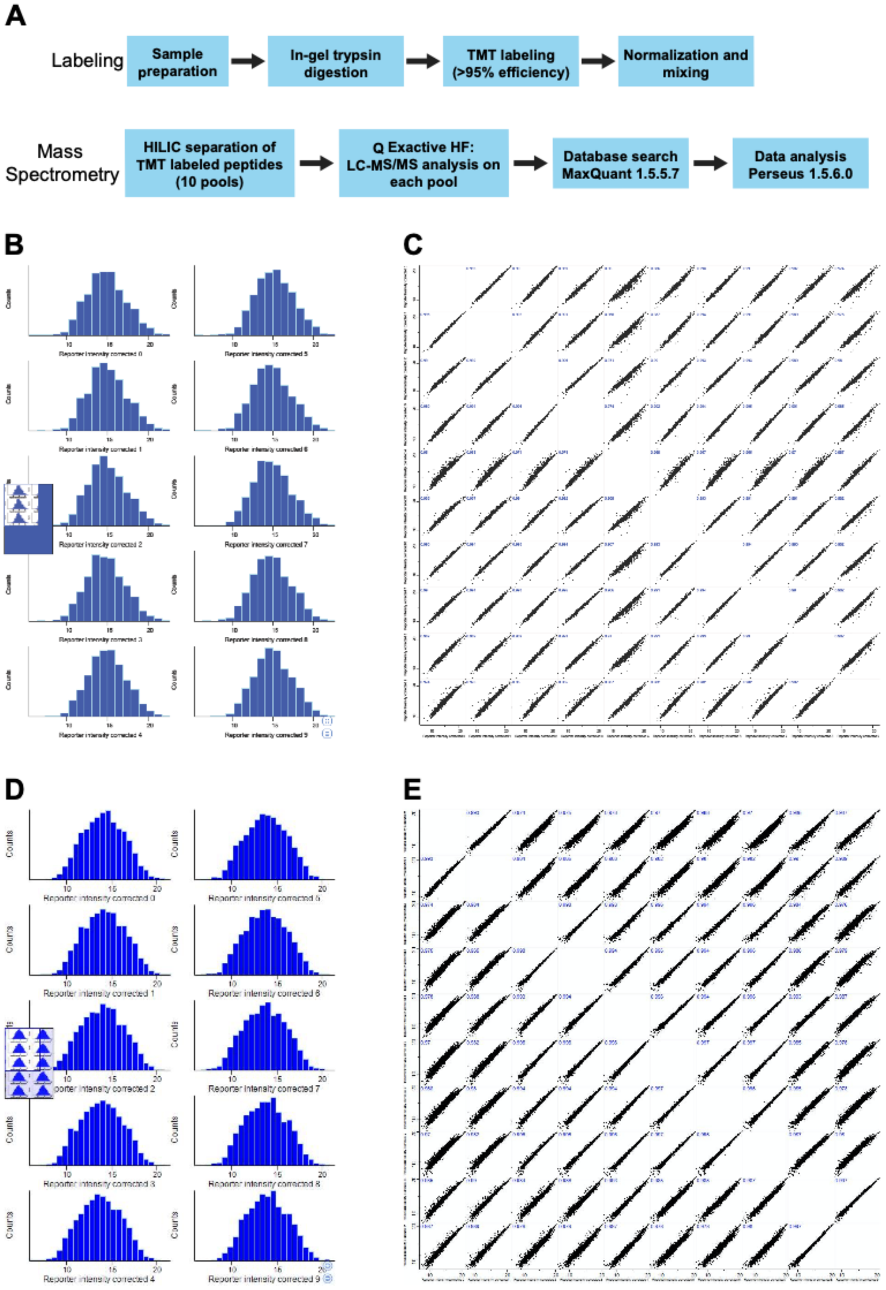

**Supplementary Figure 2. Altered synaptic composition predicts networks of functional effects in mouse models of autism.**

**A-D**) Regulatory networks for Fmr1 KO, Pten Het, and BTBR+ mice show similar upstream regulators and downstream effects. **E-I**) Regulatory networks for Cntnap2 KO, Shank3*, Cacna1c*, and Anks1b Het mice show similarities. Upstream regulators and downstream effects were predicted in IPA from fold-changes of proteins in the postsynaptic proteome.

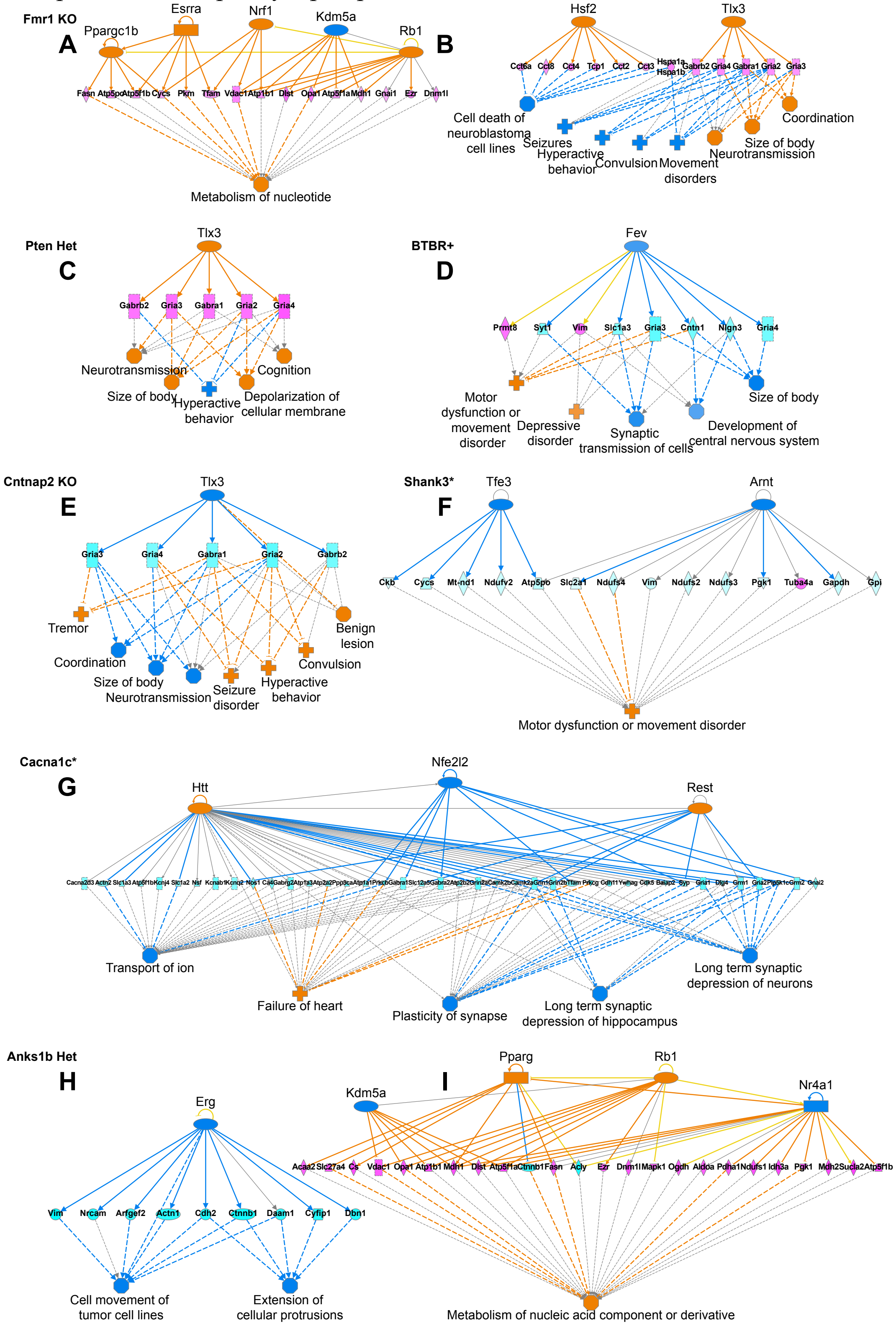

48
